# Can the lack of fibrillar form of alpha-synuclein in Lewy bodies be explained by its catalytic activity?

**DOI:** 10.1101/2021.05.09.443304

**Authors:** Ivan A. Kuznetsov, Andrey V. Kuznetsov

## Abstract

Finding the causative pathophysiological mechanisms for Parkinson’s disease (PD) is important for developing therapeutic interventions. Until recently, it was believed that Lewy bodies (LBs), the hallmark of PD, are mostly composed of alpha-synuclein (α-syn) fibrils. Recent results (Shahmoradian et al., Lewy pathology in Parkinson’s disease consists of crowded organelles and lipid membranes, Nature Neuroscience 22 (2019) 1099-1109) demonstrated that the fibrillar form of α-syn is lacking from LBs. Here we propose that this surprising observation can be explained by the catalytic activity of the fibrillar form of α-syn. We assumed that α-syn fibrils catalyze the formation of LBs, but do not become part of them. We developed a mathematical model based on this hypothesis. By using the developed model, we investigated the consequences of this hypothesis. In particular, the model suggests that the long incubation time of PD can be explained by a two-step aggregation process that leads to its development: (i) aggregation of monomeric α-syn into α-syn oligomers and fibrils and (ii) clustering of membranebound organelles, which may cause disruption of axonal trafficking and lead to neuron starvation and death. The model shows that decreasing the rate of destruction of α-syn aggregates in somatic lysosomes accelerates the formation of LBs. Another consequence of the model is the prediction that removing α-syn aggregates from the brain after the aggregation of membrane-bound organelles into LBs has started may not stop the progression of PD because LB formation is an autocatalytic process; hence, the formation of LBs will be catalyzed by aggregates of membrane-bound organelles even in the absence of α-syn aggregates. The performed sensitivity study made it possible to establish the hierarchy of model parameters with respect to their effect on the formation of vesicle aggregates in the soma.

## 1. Introduction

Parkinson’s disease (PD) is a progressive neurodegenerative disorder. No cures or treatments that slow disease progression currently exist. The only available treatments are dopaminergic drugs [1,2] and deep brain stimulation [3], which only provide symptomatic treatment. PD is caused by the death of dopaminergic (DA) neurons in the brain region called substantia nigra pars compacta (SNc). A postmortem examination found the presence of abnormal aggregates called Lewy bodies (LBs) (in the soma) and Lewy neurites (LNs) (in the axon) [3].

Until recently it was believed that misfolded fibrillar α-syn is the main component of LBs [4]. Surprisingly, a recent report found that LBs are mostly composed of membrane fragments, organelles, vesicular structures, and various lipid constituencies [5]. The reasons behind these surprising observations are still debated [6]. There is strong evidence suggesting that misfolded and fibrillated α-syn is the pathogen that causes PD [7]. The spread of misfolded α-syn occurs very slowly. It takes about 10 years for Lewy pathology (LBs and LNs) to spread into grafted neuronal cells [8,9]. Understanding whether α-syn aggregates are the PD-causing agents is important because removing the PD-causing agents should lead to remission and stabilization of the disease [10]. Thus, identifying the correct disease-causing agents can have significant impacts on developing effective disease-modifying therapies.

Despite some recent progress in modeling PD [11–13], the urgency for further progress in understanding fundamentals of PD is dictated by the need to develop disease-modifying treatments. Here we develop a model based on the hypothesis that the formation of LBs and LNs is catalyzed by α-syn aggregates (Fig. 1). This would explain why [5] did not find α-syn fibrils in LBs. Our hypothesis would also help to explain another paradox associated with PD: if misfolded α-syn is the disease-causing agent, as it is believed by many researchers [14–17], why does the premotor phase of PD take 20 years or more [3,18]? If transport of misfolded α-syn is the limiting factor, the duration of the premotor phase of the disease should be weeks rather than years [19]. We hypothesize that the long premotor phase of PD can be explained by the requirement for two consecutive aggregation processes: one that leads to the formation of pathological α-syn aggregates and a second that leads to the aggregation of various membrane-bound organelles into LBs. We hypothesize that the second aggregation process requires α-syn aggregates as a catalyst.

**Fig. 1.**
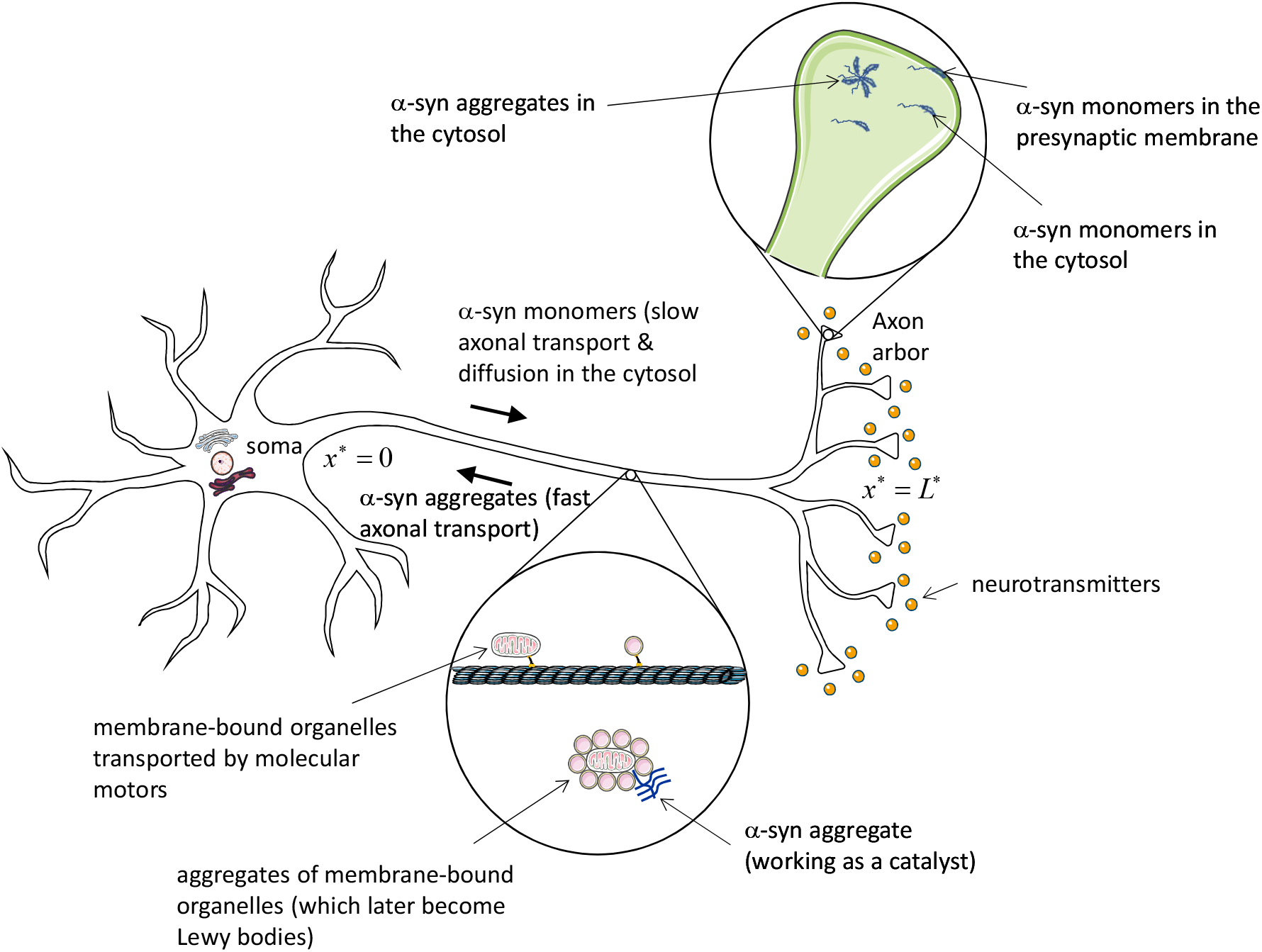
A diagram of a neuron showing transport of α-syn monomers and misfolded aggregates in the axon and catalytic effect of α-syn aggregates leading to the formation of aggregates of membrane-bound organelles (which later become LBs and LNs). Demand sites are numbered starting with the most proximal (#1) to the most distal (#*N*) [96].

## 2. Methods and models

The model is composed of five submodels detailed in sections 2.1–2.5 and illustrated in Fig. 2. In developing our model, we split all processes into two types. The first type of processes are relatively fast processes, which quickly reach steady-state and are simulated using quasi-steady-state equations. The second type of processes are extremely slow processes, which take years or decades and eventually lead to the formation of α-syn aggregates. Processes of the second-type are modeled by transient equations, as there is no steady-state for the formation of Lewy bodies. These inclusions slowly grow until a neuron dies, so their formation is an inherently transient process.

**Fig. 2.**
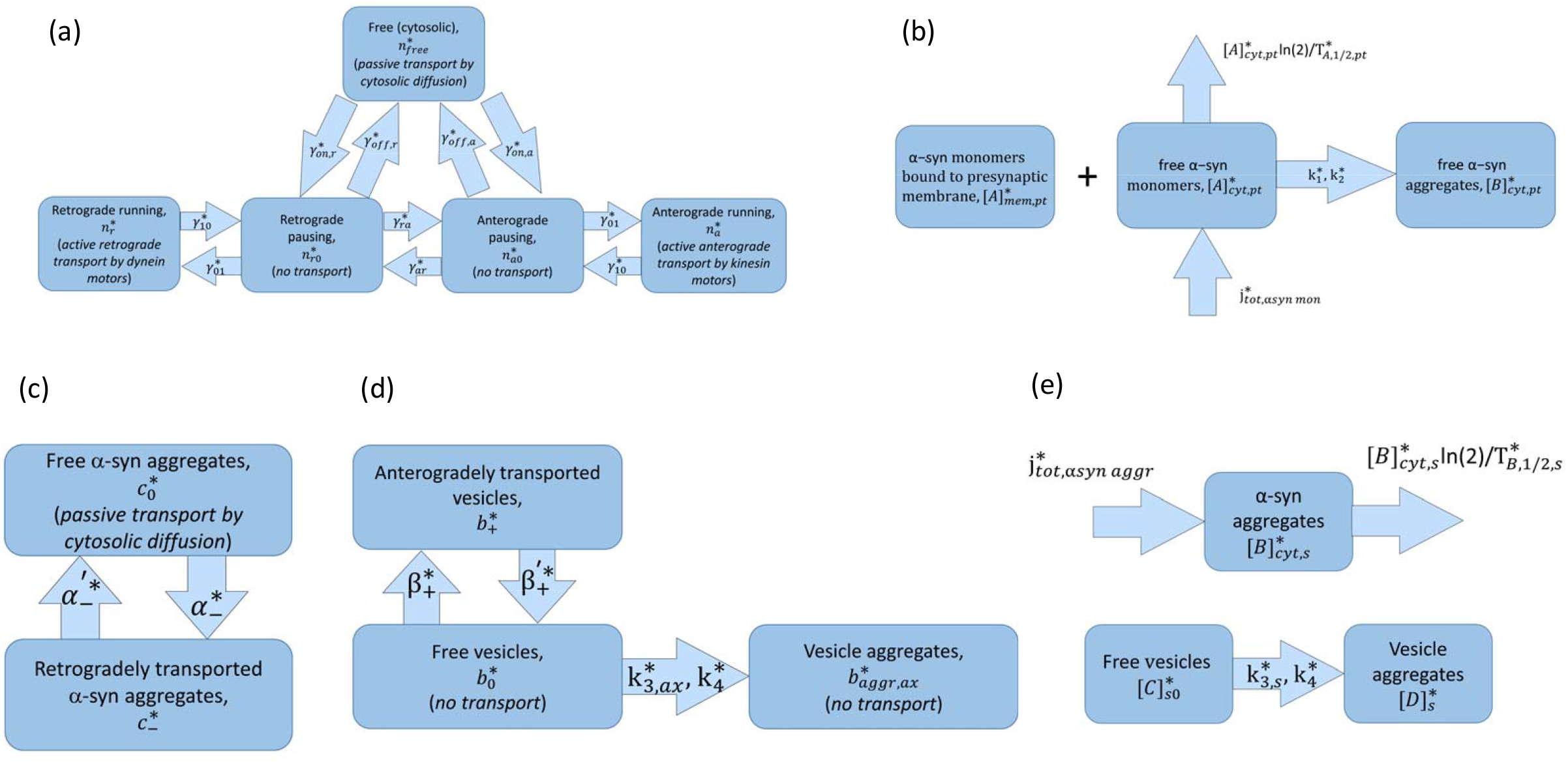
Kinetic diagrams showing various kinetic states in the following sub-models: (a) in the sub-model of slow axonal transport of α-syn monomers from the soma to the synapse. The diagram is based on the model of SCa transport of neurofilaments [25], which was modified in [26,27] to extend it to SCb transport of cytosolic proteins. (b) in the sub-model simulating transitions between various states of α-syn in the presynaptic terminal: α-syn monomers suspended in the cytosol, α-syn monomers bound to the membrane, and α-syn aggregated in the cytosol; (c) in the sub-model of fast axonal transport of α-syn aggregates from the presynaptic terminal to the soma; (d) in the sub-model of fast axonal transport of various vesicles from the soma to the presynaptic terminal and their aggregation that is presumably catalyzed by α-syn aggregates; (e) in the sub-model simulating vesicle aggregation in the soma, which is presumably catalyzed by α-syn aggregates, and also simulating the destruction of α-syn aggregates in the somatic lysosomes. The above sub-models interact with each other. For example, α-syn -monomers (Fig. 2a) enter the presynaptic terminal and form aggregates (Fig 2b). α-syn aggregates formed in the presynaptic terminal are transported to the soma (Fig. 3c). En route, the α-syn aggregates catalyze aggregation of membrane fragments from damaged vesicles that later become LNs (Fig. 2d). After exiting the axon α-syn aggregates accumulate in the soma and also catalyze aggregation of membrane fragments that later become LBs (Fig 2e). The model thus connects all the subsystems shown in Fig. 2a-e.

### 2.1. Equations simulating slow axonal transport and diffusion of α-syn monomers from the soma to the synapse (Fig. 2a)

α-syn monomers are synthesized in the soma. However, α-syn monomers have a very small concentration in the soma; they are transported to the axonal arbor, where they accumulate [20]. Transport of α-syn monomers mostly occurs in the slow component-b (SCb) [21–23]. In this paper, we neglect the 10-15% of α-syn monomers that may be transported in fast axonal transport [23]. Slow axonal transport is characterized by fast, short movements that can be either anterograde or retrograde and are probably driven by kinesin and dynein motors [24]. Rapid movements are interrupted by long pauses, during which the α-syn monomers are thought to retain some interaction with microtubules (MTs).

The sub-model suggested here for transport of α-syn in its monomeric form is based on the model of slow axonal transport of neurofilaments (transported in slow component-a, SCa). This model was developed in [25]. In our previous work, we extended this model to cytosolic proteins, which are transported in SCb and can diffuse in the cytosol [26,27]. For the α-syn monomers, the model contains anterograde and retrograde motor-driven states with concentrations 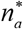 and 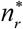, respectively (Fig. 2a). The following equations are obtained by stating the conservation of α-syn in these motor-driven states:

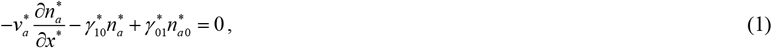

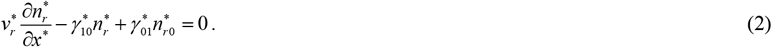

The first terms on the left-hand sides of Eqs. (1) and (2) describe anterograde (occurring with a velocity 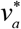) and retrograde (occurring with a velocity 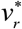) motions of the α-syn monomers, respectively, while other terms describe transitions between motor-driven and pausing kinetic states. Various kinetic states and transitions between them are depicted in Fig. 2a; the transitions are characterized by kinetic rates *γ*\*s. Asterisks denote dimensional quantities.

Equations for the α-syn monomers in two pausing states, with concentrations 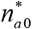 and 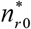, are also included in the model (Fig. 2a). Despite pausing, it is believed that the α-syn monomers retain some interaction with the MTs and can resume anterograde or retrograde motion. Only the terms that characterize transitions to/from other kinetic states are used in equations expressing conservation of α-syn monomers in the pausing states.

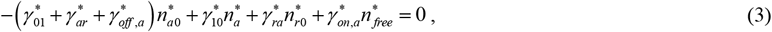

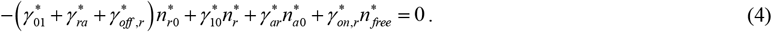

α-syn monomers in the free (cytosolic) state can be transported by diffusion. Monomeric α-syn can be also degraded in proteasomes [28], which are present in various locations in the axon [29]. It should be noted that α-syn monomers undergoing slow axonal transport in the axon may be protected from significant degradation while in transit, which may result in their longer half-life during axonal transport [30,31]. Cytosolic α-syn monomers can also transition to anterograde or retrograde pausing states. The following equation is obtained by stating the conservation of free monomeric α-syn:

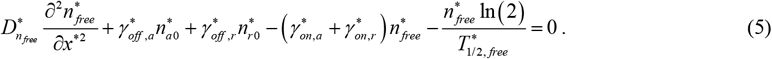

The following boundary conditions are utilized for Eqs. (1)–(5):

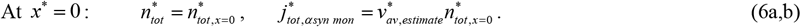

Eq. (6a) postulates the concentration of free (cytosolic) α-syn monomers at the axon entrance while Eq. (6b) postulates the total flux of newly synthesized α-syn monomers that enter the axon.

The flux of α-syn monomers (positive if anterograde) is found from the following equation:

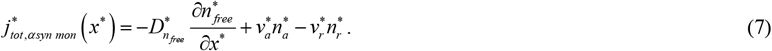

The total concentration of α-syn monomers in all five kinetic states displayed in Fig. 2a is found from the following equation:

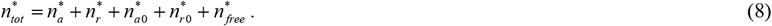

The following boundary conditions were utilized at the axon presynaptic terminal:

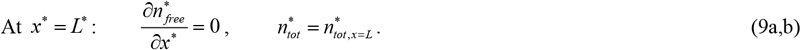

### 2.2. Equations simulating transitions between cytoplasmic and membrane-bound α-syn monomers in the presynaptic terminal, clearance of cytoplasmic α-syn monomers in proteasomes, and aggregation of cytoplasmic α-syn monomers (Fig. 2b)

Although α-syn is an intrinsically unstructured protein, it can also interact with membranes [43]. For membrane-bound α-syn, diffusion is restricted to two dimensions, which increases the likelihood of interaction between α-syn monomers and the formation of aggregates [44,45].

In this paper, we simulate aggregation of α-syn monomers (and aggregation of membrane-bound vesicles) using the minimalistic 2-step Finke-Watzky (F-W) model. The first step in the F-W model describes nucleation, *A* → *B*, and is characterized by the rate constant 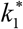. The second step describes aggregate autocatalytic growth, *A*+*B* → 2*B*, and is characterized by the rate constant 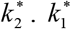 and 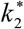 represent effective rates, hiding such processes as fragmentation [46]. In these equations, *A* represents a precursor protein and *B* represents an average of different aggregates of various sizes, including dimers and early oligomers [32,47,48].

We simulate the cytosol of the presynaptic terminal and the presynaptic membrane as two compartments that are in equilibrium. The equation expressing conservation of α-syn monomers (Fig. 1) in the cytosol of the presynaptic terminal is

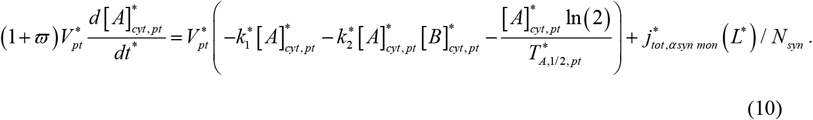

The equation for the concentration of α-syn monomers in the presynaptic membrane is

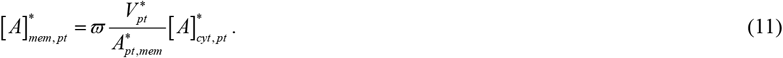

The equation simulating the production of α-syn aggregates (see Fig. 1) in the cytosol of the presynaptic terminal is

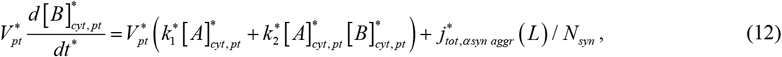

where the flux of α-syn aggregates leaving the presynaptic terminal is

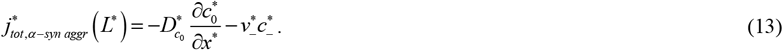

Due to the large number of synapses in DA neuron arbors, in the last terms on the right-hand side of Eqs. (10) and (12) we divided the flux of α-syn monomers coming through the axon to the synapse and the flux of α-syn aggregates transported from the whole arbor to the soma by the number of synapses.

While aggregation of α-syn occurs in the cytosol, nucleation of α-syn aggregates is more likely to occur when α-syn is associated with a cellular membrane [45,49]. For that reason, we assumed that the rate constant for the first step of the F-W model for aggregation of α-syn monomers in the cytosol of the presynaptic terminal, 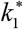, is proportional to the area of the active zone of the presynaptic membrane, which is involved in the release of neurotransmitters:

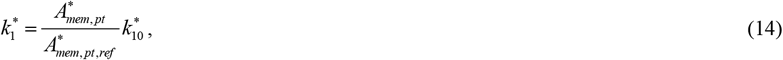

where 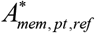 is a reference area of the active zone of the presynaptic membrane.

For estimating the volume of the presynaptic terminal, we approximated the terminal as a hemisphere with a diameter 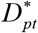. The area of the presynaptic membrane, which is involved in the release of neurotransmitters from the presynaptic terminal, is then estimated as that of a circle with the diameter 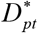:

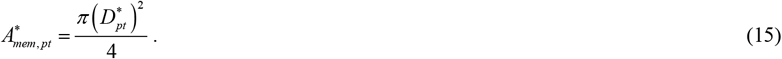

The flux of the α-syn monomers in the axon is calculated as:

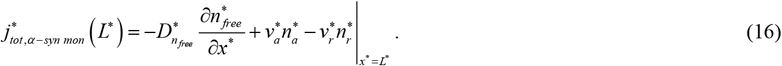

The following initial conditions are used for Eqs. (10) and (12):

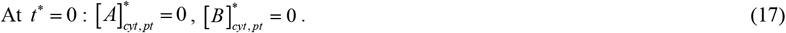

Eq. (17) assumes that initially, the presynaptic terminal does not contain any α-syn monomers, which must be first produced in the soma and then transported to the presynaptic terminal. This is a simplifying assumption because to accurately model the development of α-syn concentration in the presynaptic terminal toward steady-state the growth of the axon needs to be simulated.

### 2.3. Equations simulating fast axonal transport of α-syn aggregates inside autophagic vesicles from the presynaptic terminal to the soma (Fig. 2c)

α-syn aggregates are believed to be cleared in lysosomes located in the soma [56]. α-syn aggregates are delivered to lysosomes by autophagy [57,58]. Autophagosomes move in the fast retrograde component [59], propelled by dynein motors [60,61].

We thus assume that α-syn aggregates are transported retrogradely from the presynaptic terminal to the soma in autophagic vesicles. In the soma, α-syn aggregates are destroyed in lysosomes. Since we consider vesicles that undergo fast axonal transport, in this sub-model we assume that α-syn aggregates can be in two kinetic states: retrograde, when they are transported inside vesicles by retrograde motors, and free, when they are suspended in the cytosol (Fig. 2c). Applying conservation of retrogradely transported α-syn aggregates results in the following equation:

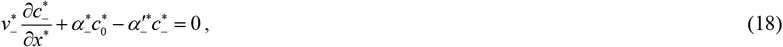

where the first term on the left-hand side describes retrograde motor-driven transport of α-syn aggregates and the last two terms describe the transition of α-syn aggregates to and from the free cytosolic state (Fig. 2c). 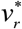 in Eq. (2) and 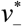 in Eq. (18) characterize the retrograde velocities of α-syn monomers and α-syn aggregates, respectively, due to the action of dynein motors. However, these are not velocities of dynein motors per se, but rather effective parameters. This is because modes of axonal transport (slow versus fast), the average size of the cargo, the average number of motors transporting the cargo, and other parameters that characterize the transport of α-syn monomers and aggregates are different. Therefore, 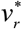 and 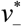 may take on different values.

The presence of clusters of fragmented membranous material in LBs [5] suggests that autophagosomes may get damaged during their retrograde transport [62,63], and misfolded α-syn aggregates may escape to the cytosol. Conservation of free α-syn aggregates that are detached from MTs is expressed by the following equation:

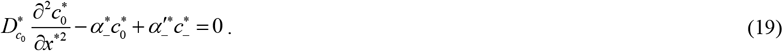

Eqs. (18) and (19) are solved subject to the following boundary conditions. At the axon hillock, we impose the following condition:

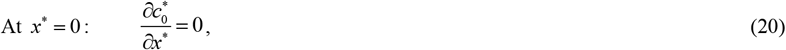

where the flux of α-syn aggregates can be found from the following equation:

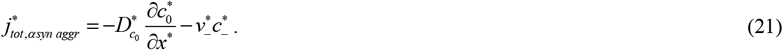

At the axon presynaptic terminal, we imposed the following boundary conditions:

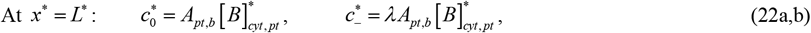

where

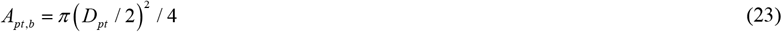

is the cross-sectional area of the branch connected to the presynaptic terminal. We assumed that the diameter of the branch is half of the diameter of the presynaptic terminal since proteins and vesicles should be able to pass through it.

Eq. (22a) gives the concentration of free (cytosolic) α-syn aggregates in the presynaptic terminal and Eq. (22b) gives the concentration of α-syn aggregates entering the axon from the presynaptic terminal moving retrogradely, which are propelled by dynein motors.

### 2.4. Equations simulating anterograde fast axonal transport of membrane (via anterograde transport of various membrane-bound vesicles) from the soma to the presynaptic terminal and catalysis of formation of vesicle aggregates (which eventually form LNs) by aggregated α-syn in the axon (Fig. 2d)

We assume that various membrane-bound vesicles synthesized in the soma are transported toward the presynaptic terminal anterogradely by fast axonal transport. This leads to the flow of the membrane from the soma toward the synapse. Since vesicles are transported by kinesin motors pulling them along MTs, one of the kinetic states in the model represents anterogradely moving vesicles. It is also assumed that vesicles can detach from MTs and become freely suspended in the cytosol (Fig. 2d). Applying conservation of anterogradely transported membrane gives the following equation:

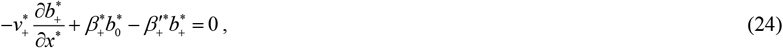

where the first term on the left-hand side describes anterograde motor-driven vesicle transport and the last two terms describe transitions between anterograde motor-driven state and free (cytosolic) state (Fig. 2d). 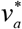 and 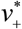 characterize the anterograde velocities of α-syn monomers and anterogradely transported vesicles, respectively, due to the action of kinesin motors. These, however, are not kinesin motor velocities but rather effective parameters. This is due to the differences in axonal transport modes (slow versus fast), cargo average size, the average number of motors interacting with cargo, and other parameters distinguishing α-syn and vesicles transport. As a result, values of 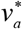 and 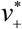 may be different.

Conservation of membrane in the vesicles that are detached from MTs and freely suspended in the cytosol is expressed by the following equation:

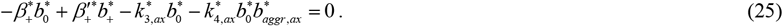

The first two terms in Eq. (25) are kinetic terms describing transitions between anterogradely running and free kinetic states of vesicles (Fig. 2c). The last two terms in Eq. (25) describe the decrease in the concentration of free vesicles in the cytosol due to the production of vesicle aggregates. Simulation of this process is important for understanding the formation of LBs, which involves the sequestration of proteins, organelles, lipids, and endomembranes and their packing and compaction into LB-like inclusions [5,66,67]. The last two terms in Eq. (25) are written utilizing the F-W model. The mechanism of vesicle aggregates formation combines primary nucleation (described by the kinetic constant 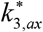 in Eq. (25)) and secondary nucleation (described by the kinetic constant 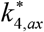). Accumulation of membrane contained in vesicle aggregates is described by the following equation:

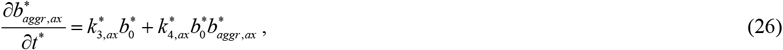

where the diffusivity of vesicle aggregates was neglected because of their large size.

We assumed that the nucleation step in the formation of vesicle aggregates is catalyzed by α-syn aggregates, for which reason α-syn aggregates do not become a part of LBs. It was observed that α-syn fibrils can “absorb” lipid vesicles causing their clustering [68]. Therefore, it is possible that while misfolded α-syn aggregates are delivered to the soma for degradation, they catalyze clustering of various membrane-bound vesicles in the axon. Thus, we assume that the rate of formation of vesicle aggregates in the axon is proportional to the concentration of free (suspended in the cytosol) α-syn aggregates in the axon:

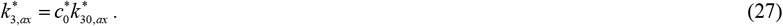

Eqs. (24) and (25) must be solved subject to the following boundary condition:

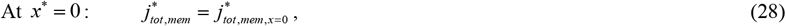

where

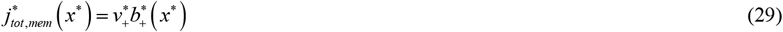

and 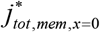 is the rate at which membrane contained in membrane-bound vesicles enters the axon from the soma (the rate of synthesis of the membrane contained in the axonal cargo in the soma).

Eq. (26) must be solved subject to the following initial condition:

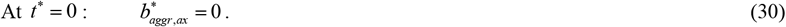

### 2.5. Equations describing the destruction of α-syn aggregates in the somatic lysosomes and catalysis of the formation of vesicle aggregates (which ultimately leads to the formation of LBs) by α-syn aggregates in the soma (Fig. 2e)

It is likely that under normal conditions α-syn monomers are destroyed in proteasomes while α-syn aggregates are destroyed in lysosomes [28]. We thus assumed that α-syn monomers are degraded in proteasomes, which are present in various locations in the axon, while α-syn aggregates are degraded in lysosomes, which are mostly located in the soma [71].

The process of destruction of α-syn aggregates in the lysosomes in the soma is modeled by the following equation:

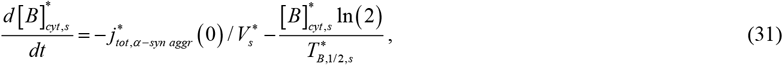

where

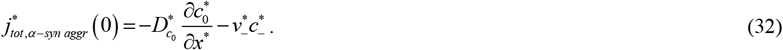

The negative sign before the first term of the right-hand side of Eq. (31) is because α-syn aggregates enter the soma by retrograde transport from the axon.

We assumed that α-syn aggregates catalyze the formation of vesicle aggregates in the soma. We used the F-W model to simulate the formation of vesicle aggregates:

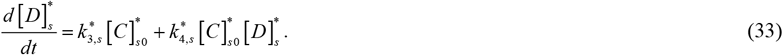

Similar to Eq. (27), we assumed that the nucleation step in the formation of vesicle aggregates is catalyzed by α-syn aggregates, and thus the rate of formation of vesicle aggregates in the soma is proportional to the concentration of α-syn aggregates in the soma:

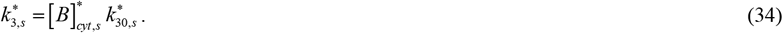

Eqs. (31), (33), and (34) must be solved subject to the following initial conditions:

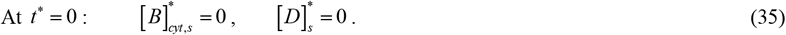

## 3. Results

### 3.1. Analysis of various concentrations and fluxes predicted by the model

The model predicts that most α-syn monomers localize in the presynaptic terminal, which is consistent with [43,74] (Fig. 3a). The flux of α-syn monomers along the axon is uniform because we assumed that α-syn monomers are protected from degradation during their transport in the axon, and hence under steady-state conditions, their flux must be independent of the position in the axon (Fig. 3b).

**Fig. 3.**
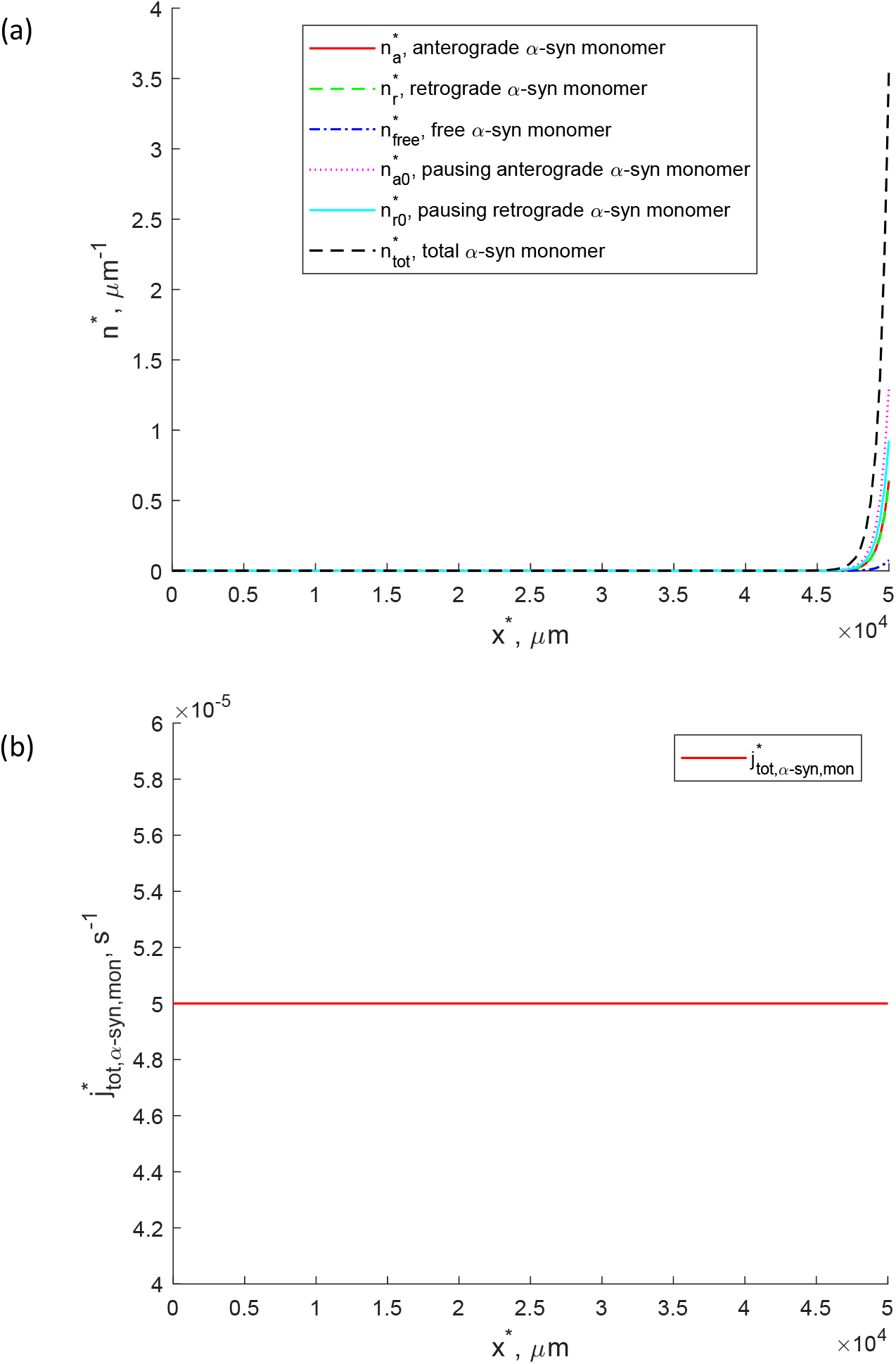
(a) Various linear concentrations (per unit length of the axon) of α-syn monomers in the axon, including the total concentration, which is the sum of α-syn concentrations in motor-driven, pausing, and diffusing states. (b) Total flux of α-syn monomers due to active transport, powered by molecular motors, and diffusion in the cytosol. 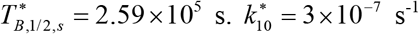.

α-syn monomers exist in the cytosol of the presynaptic terminal in equilibrium with membrane-bound α-syn monomers [43]. Both volumetric and surface concentrations quickly reach their steady-state values and after that remain stationary (Fig. 4a). The concentration of α-syn aggregates in the cytosol also quickly reaches its steady-state value and after that remains stationary (Fig. 4b). This equilibrium value is controlled by the rate of production of aggregates in the presynaptic terminal and their removal into the axon by retrograde transport to the soma for degradation in lysosomes.

**Fig. 4.**
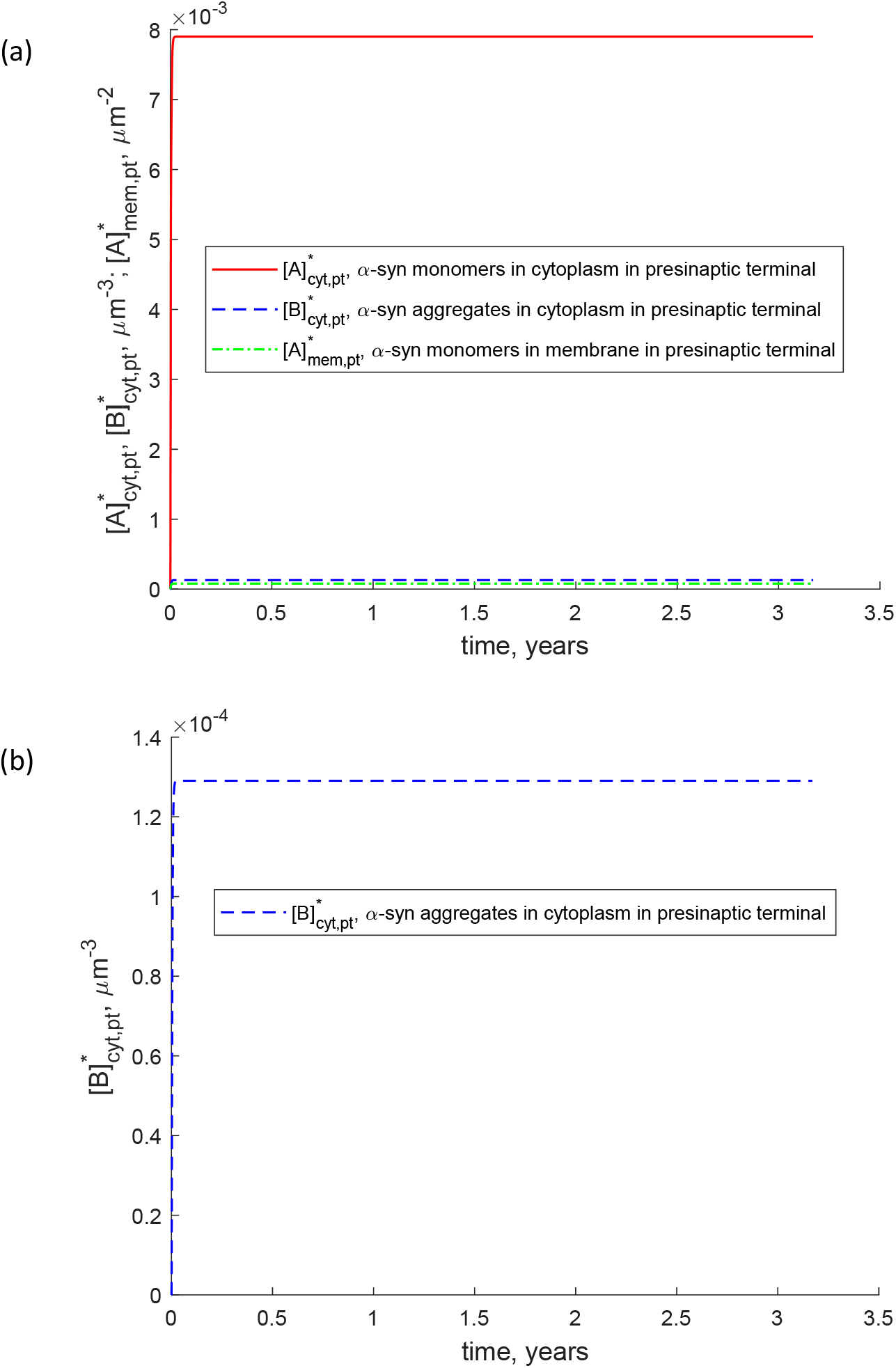
(a) Volumetric concentrations of α-syn monomers and α-syn aggregates in the presynaptic terminal and surface concentration of α-syn monomers bound to the presynaptic membrane. (b) Volumetric concentration of α-syn aggregates in the presynaptic terminal (same as in (a), but on the enlarged scale). 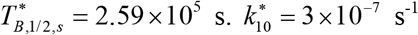.

Linear number densities of retrogradely transported α-syn aggregates and free α-syn aggregates in the axon are uniform along the axon (Fig. 5). They take on the same values due to the choice of values of kinetic constants, 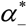 and 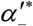 (Table 7).

**Fig. 5.**
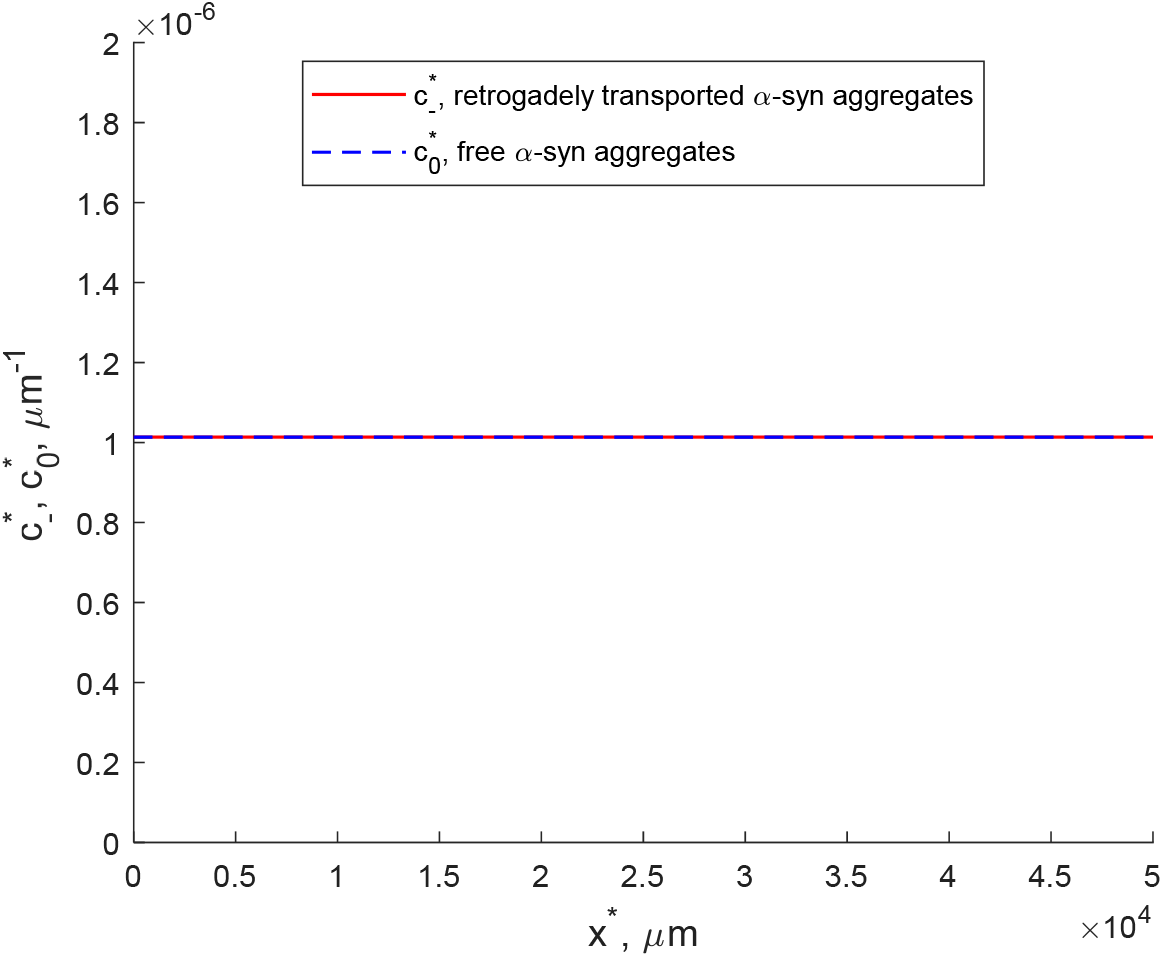
Linear concentrations (per unit length of the axon) of retrogradely transported α-syn aggregates and free α-syn aggregates in the axon. 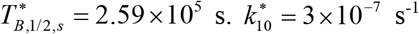.

**Table 1.** Independent variables used in the mathematical model of α-syn monomer transport in the axon.

| Symbol | Definition | Units |
| --- | --- | --- |
| $x^*$ | Cartesian coordinate that originates at the axon hillock and is directed along the axon toward the presynaptic terminal (Fig. 1) | $\mu\text{m}$ |
| $t^*$ | Time | s |

**Table 2.**
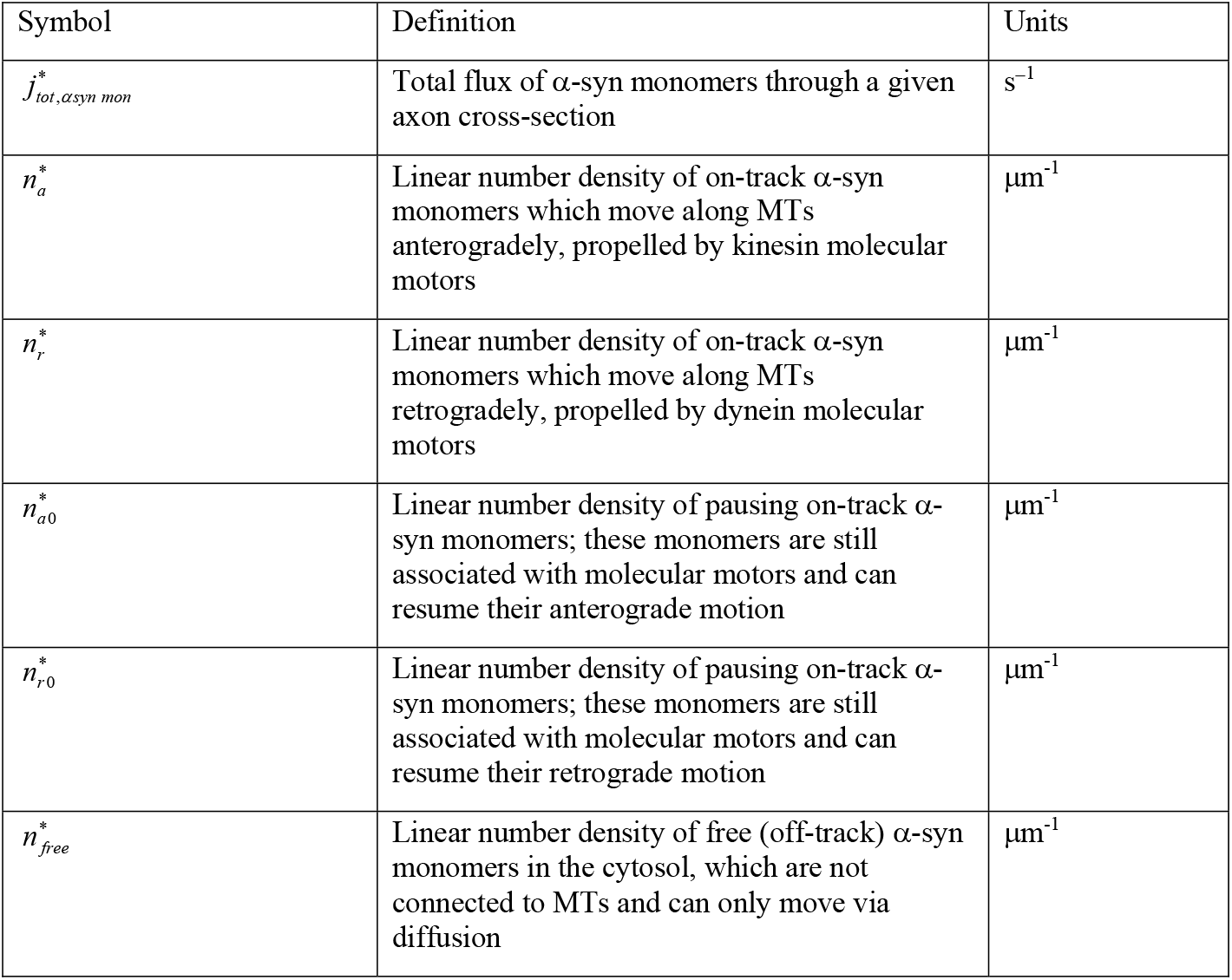
Dependent variables used in the mathematical model of α-syn monomer transport in the axon.

**Table 3.**
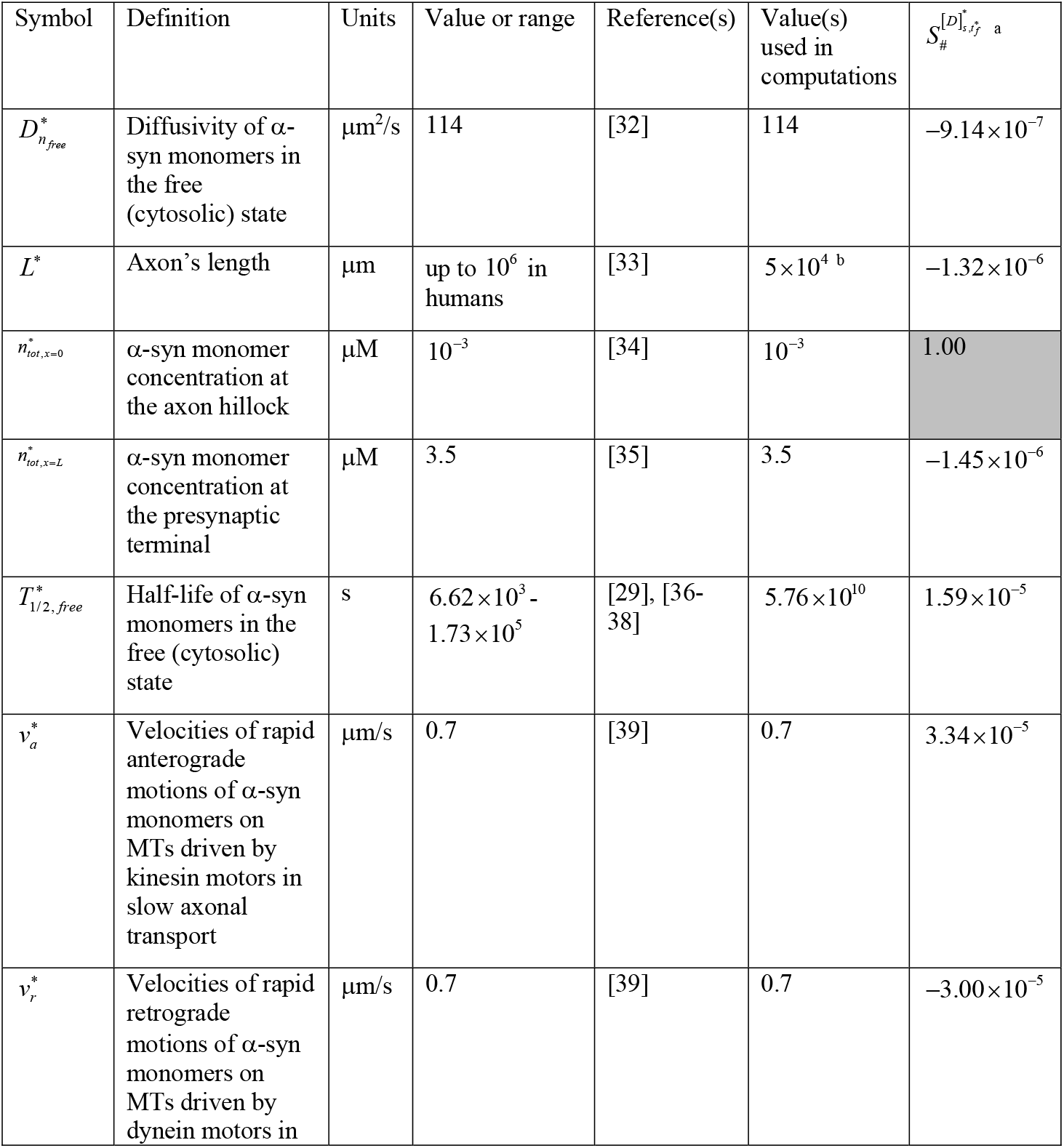
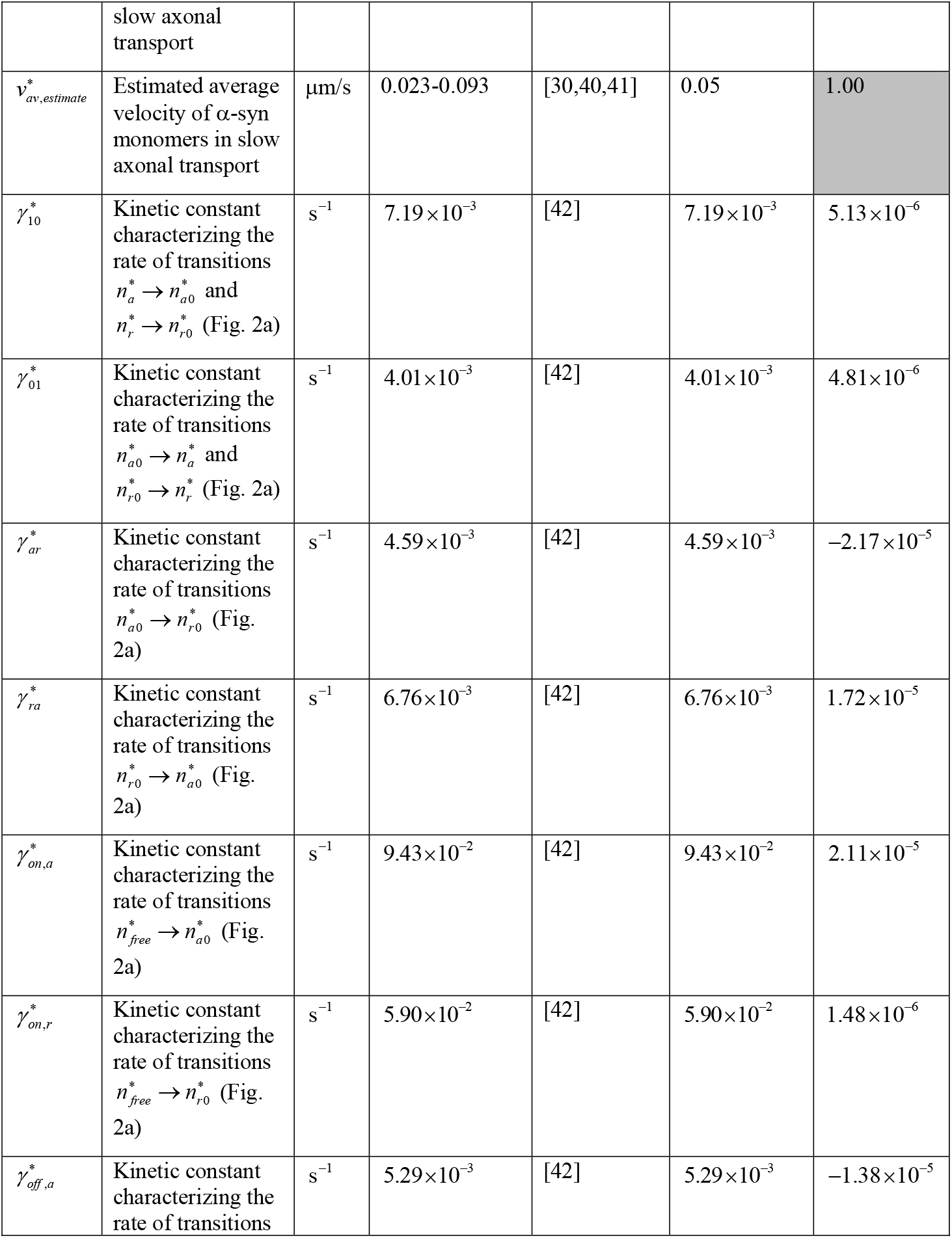
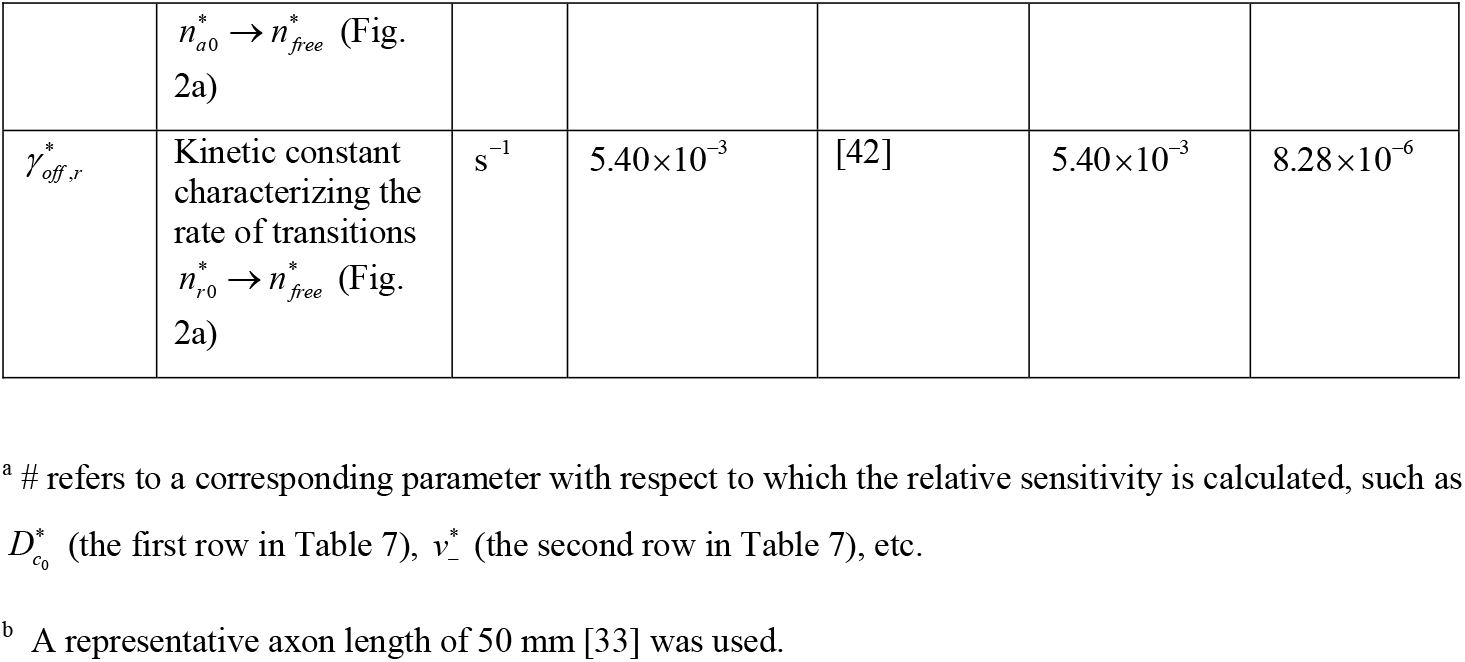
Parameters used to characterize the transport of α-syn monomers in the axon. The sensitivity of the area of membrane contained in vesicle aggregates in the soma to these parameters (see section 3.2) is given in the last column. The cells with the largest sensitivity coefficients are shaded grey.

**Table 4.** Dependent variables in the model of α-syn monomers accumulation in the cytosol and membrane of the presynaptic terminal as well as production of α-syn aggregates in the cytosol of the presynaptic terminal.

| Symbol | Definition | Units |
| --- | --- | --- |
| $[A]_{cyt,pt}^*$ | Volumetric concentration of α-syn monomers in the cytosol of the presynaptic terminal | $\mu\text{m}^{-3}$ |
| $[A]_{mem,pt}^*$ | Surface concentration of α-syn monomers bound to the presynaptic membrane | $\mu\text{m}^{-2}$ |
| $[B]_{cyt,pt}^*$ | Volumetric concentration of α-syn aggregates in the cytosol of the presynaptic terminal | $\mu\text{m}^{-3}$ |

**Table 5.** Parameters characterizing accumulation of α-syn monomers in the cytosol and membrane in the presynaptic terminal as well as production of α-syn aggregates in the cytosol of the presynaptic terminal. The sensitivity of the area of membrane contained in vesicle aggregates in the soma to these parameters (see section 3.2) is given in the last column. The cells with the largest sensitivity coefficients are shaded grey.

| Symbol | Definition | Units | Value or range | Reference(s) | Value(s) used in computations | $S_{\#}^{[D]_{s,tf}^*}$ |
| --- | --- | --- | --- | --- | --- | --- |
| $A_{mem,pt}^*$ | Area of the active zone of the presynaptic membrane, which is involved in the release of neurotransmitters from the presynaptic terminal | $\mu\text{m}^2$ | $3.14 \times 10^{-2}$ - $1.39 \times 10^{-1}$ <sup>a</sup> | [50,51] | $3.14 \times 10^{-2}$ | $9.76 \times 10^{-1}$ |
| $A_{mem,pt,ref}^*$ | Reference area of the active zone of presynaptic membrane | $\mu\text{m}^2$ | | | $3.14 \times 10^{-2}$ <sup>b</sup> | $-9.75 \times 10^{-1}$ |
| $k_{10}^*$ | Rate constant for the first step of the F-W model for aggregation of α-syn monomers in the cytosol of the presynaptic terminal | $\text{s}^{-1}$ | $2.78 \times 10^{-7}$ - $9.44 \times 10^{-5}$ | [32] | $3 \times 10^{-7}$, $3 \times 10^{-10}$ | $9.76 \times 10^{-1}$ |
| $k_2^* N_A^*$ | Rate constant for the second step of the F-W model for aggregation of α-syn monomers in the cytosol of the presynaptic terminal | $\mu\text{m}^3 \text{s}^{-1} \text{mol}^{-1}$ | $1.19 \times 10^{15}$ - $5.00 \times 10^{15}$ | [32] | $2 \times 10^{15}$ | $1.39 \times 10^{-6}$<br>($S_{k_2^*}^{[D]_{s,tf}^*}$) |
| $N_A^*$ | Avogadro constant | $\text{mol}^{-1}$ | $6.022 \times 10^{23}$ | | $6.022 \times 10^{23}$ | |
| $N_{syn}$ | Number of synapses in a dopamine neuron | | $2.45 \times 10^5$ - $2.4 \times 10^6$ | [52,53] | $2.45 \times 10^5$ | $-2.33 \times 10^{-3}$ |
| $T_{A,1/2,pt}^*$ | Half-life of α-syn in the monomeric form in the cytosol of the presynaptic terminal | s | $6.62 \times 10^3$ - $1.73 \times 10^5$ | [29,36-38] | $5.76 \times 10^4$ | $9.72 \times 10^{-1}$ |
| $V_{pt}^*$ | Volume of the presynaptic terminal | $\mu\text{m}^3$ | $2.09 \times 10^{-3} - 1.94 \times 10^{-2}$ | [50,51]<sup>a</sup> | $2.09 \times 10^{-3}$ | $-2.33 \times 10^{-3}$ |
| $\varpi$ | The ratio of the number of α-syn molecules bound to the presynaptic membrane to that in the cytosolic state in the presynaptic terminal | | 0.1-0.15 | [54,55] | 0.15 | $-5.20 \times 10^{-4}$ |

<sup>a</sup> The diameter of the presynaptic terminal is estimated to be between 0.20 $\mu\text{m}$ [50] and 0.42 $\mu\text{m}$ [51].
<sup>b</sup> We used the same value as the area of the presynaptic terminal.

**Table 6.**
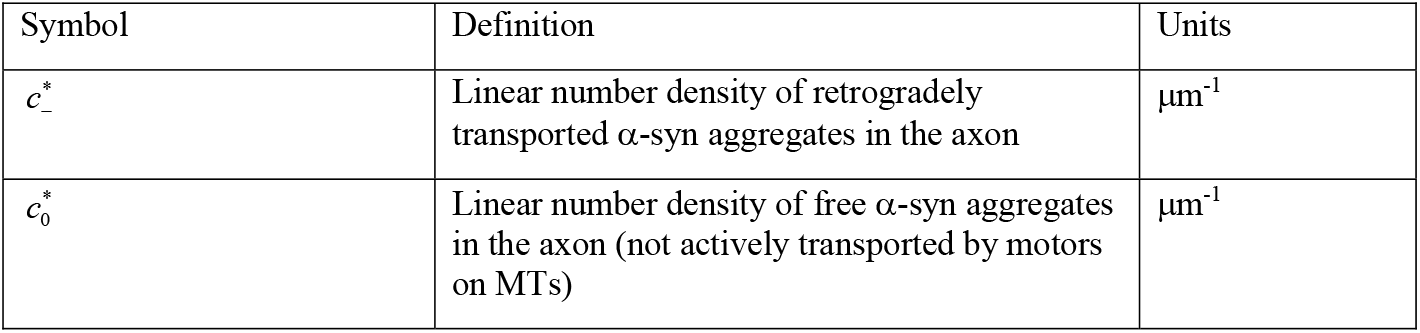
Dependent variables in the model of retrograde transport of α-syn aggregates.

| Symbol | Definition | Units |
| --- | --- | --- |
| $c_-^*$ | Linear number density of retrogradely transported α-syn aggregates in the axon | $\mu\text{m}^{-1}$ |
| $c_0^*$ | Linear number density of free α-syn aggregates in the axon (not actively transported by motors on MTs) | $\mu\text{m}^{-1}$ |

**Table 7.** Parameters characterizing retrograde transport of α-syn aggregates and their estimated values. The sensitivity of the area of membrane contained in vesicle aggregates in the soma to these parameters (see section 3.2) is given in the last column.

| Symbol | Definition | Units | Value or range | Reference(s) | Value(s) used in computations | $S_{\#}^{[D]_{s,j_f}^*}$ |
| --- | --- | --- | --- | --- | --- | --- |
| $D_{c_0}^*$ | Diffusivity of α-syn aggregates in the cytosolic state | $\mu\text{m}^2/\text{s}$ | 78 | [32] | 78 | $7.29 \times 10^{-13}$ |
| $v_-^*$ | Retrograde velocity of α-syn aggregates which are transported in vesicles propelled by dynein motors | $\mu\text{m}/\text{s}$ | 1.2-1.3 | [64] | 1.2 | $2.33 \times 10^{-3}$ |
| $\alpha_-^*$ | Kinetic constant characterizing the rate of transition $c_0^* \rightarrow c_-^*$ (Fig. 2c) | $\text{s}^{-1}$ | 0.1-1 | [65] | 1 | $2.29 \times 10^{-3}$ |
| $\alpha_-'^*$ | Kinetic constant characterizing the rate of transition $c_-^* \rightarrow c_0^*$ (Fig. 2c) | $\text{s}^{-1}$ | 0.1-1 | [65] | 1 | $-2.29 \times 10^{-3}$ |
| $\lambda$ | Degree of loading of α-syn aggregates that characterizes association of these aggregates with dynein motors at the presynaptic terminal | | 0.1-10 | [65] | 1 | $4.15 \times 10^{-5}$ |

The total area of membrane contained in all membrane-bound organelles traveling anterogradely in the axon and suspended in the axon’s cytoplasm is uniform along the axon (Fig. 6). The flux of anterogradely transported vesicles in the axon is also uniform due to a small rate of vesicle aggregation (small values of 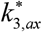 and 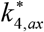, Table 9 and Eq. (27)).

**Fig. 6.**
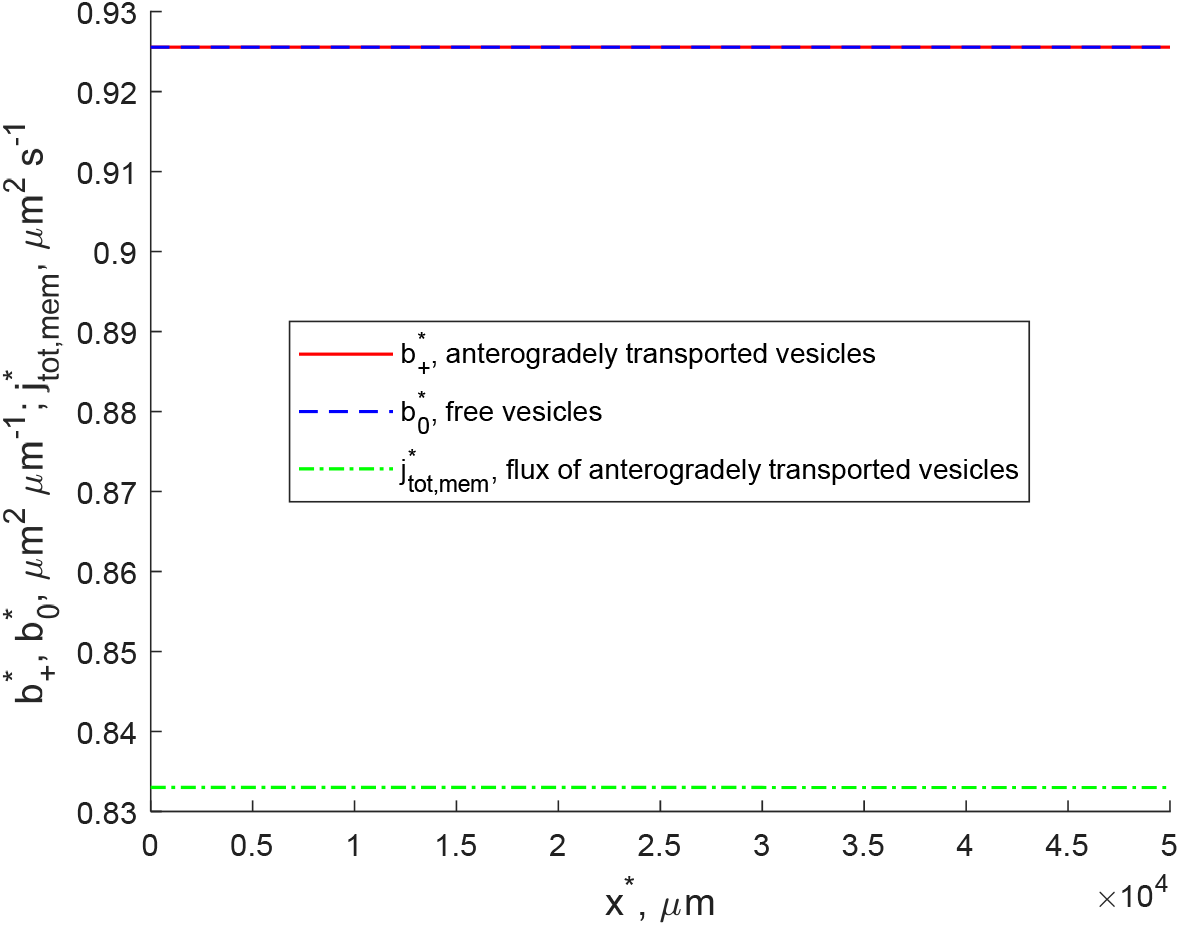
Total area of membrane in all membrane-bound organelles that are actively anterogradely transported in the axon and in all membrane-bound organelles that are detached from MTs, per unit length of the axon. Also, the anterograde flux of membrane due to transport of membrane-bound organelles. 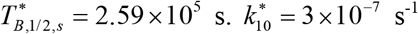.

**Table 8.**
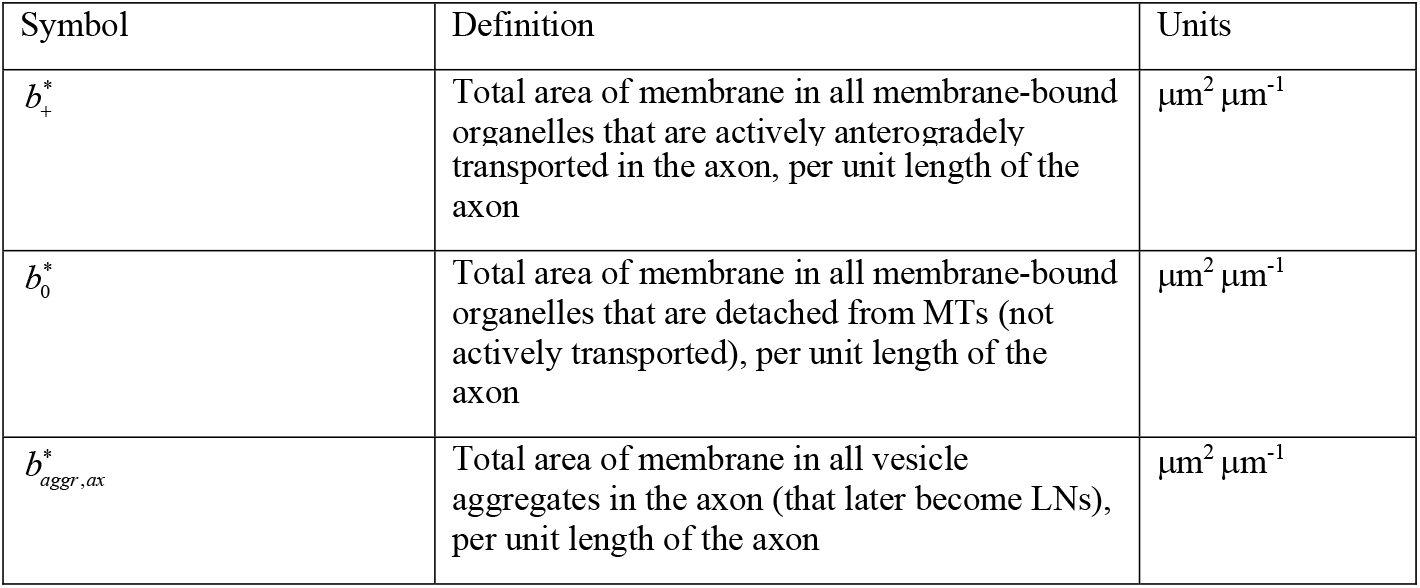
Dependent variables in the model of anterograde transport of membrane in synaptic vesicles and its aggregation in the axon.

**Table 9.**
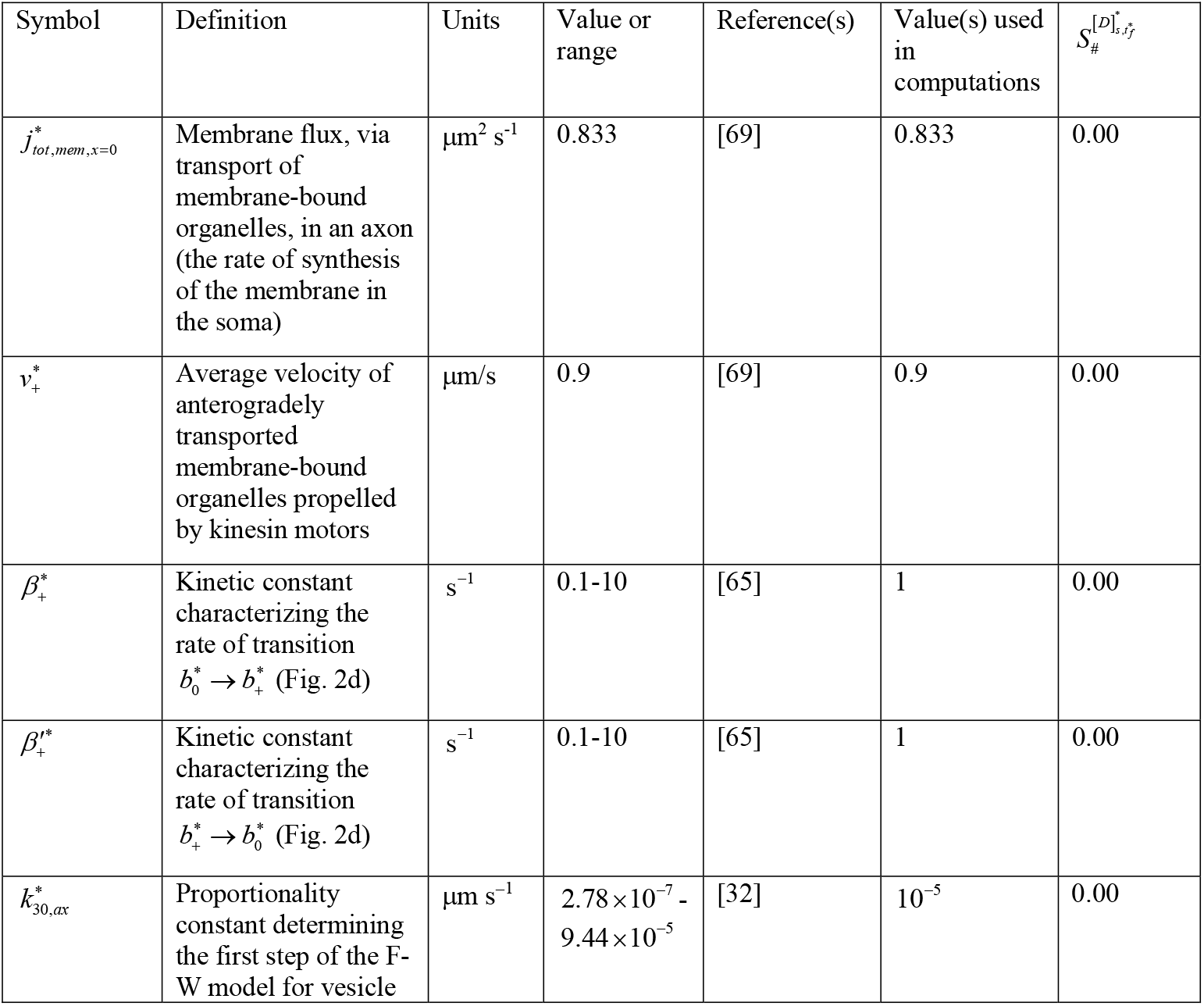
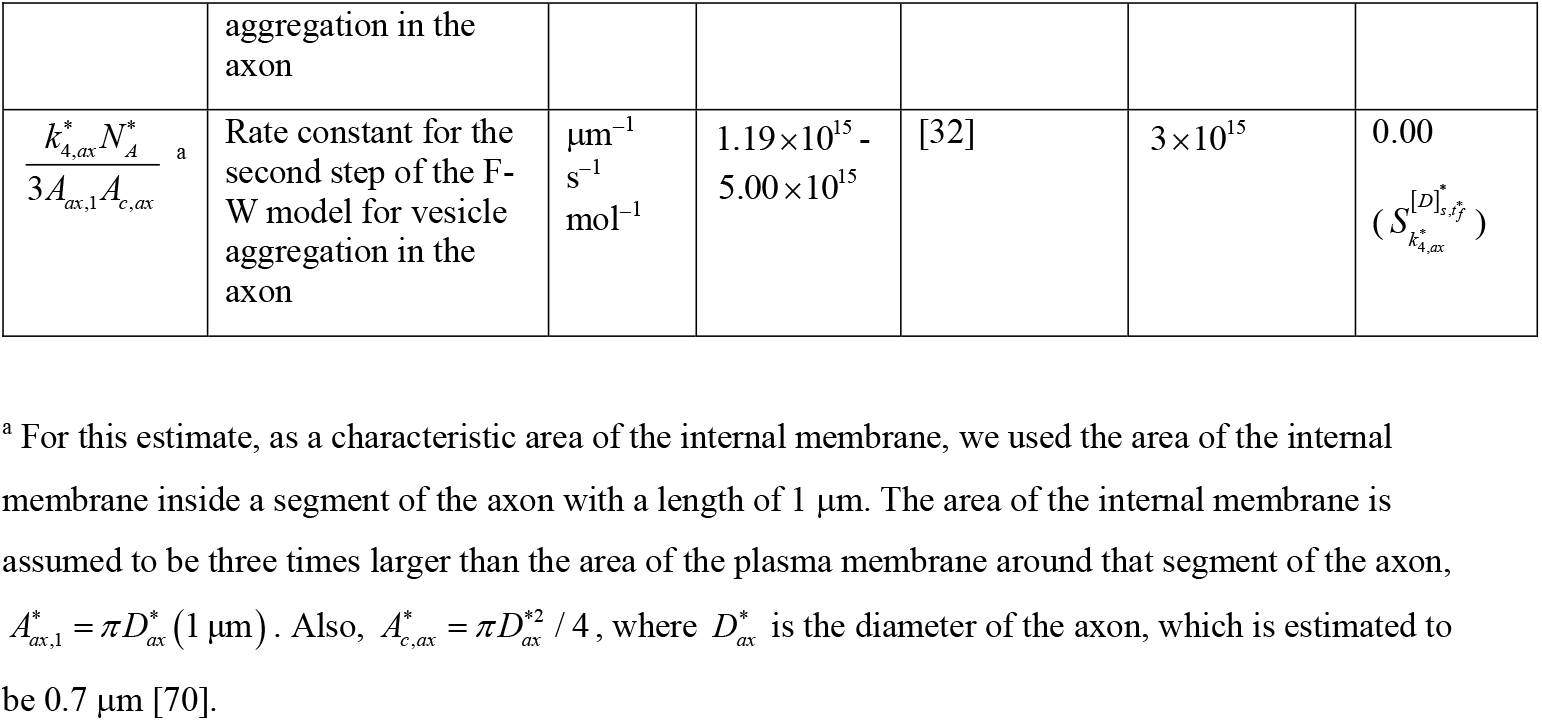
Parameters characterizing anterograde vesicle transport and aggregation in the axon. The sensitivity of the area of membrane contained in vesicle aggregates in the soma to these parameters (see section 3.2) is given in the last column.

The total area of the membrane in all vesicle aggregates in the axon is uniformly distributed along the length of the axon (Fig. 7a). The total area of the membrane in all vesicle aggregates in the middle of the axon monotonically (but not linearly) increases with time, with the rate of the increase becoming larger over time. This follows from the curvature of the line in Fig. 7b.

**Fig. 7.**
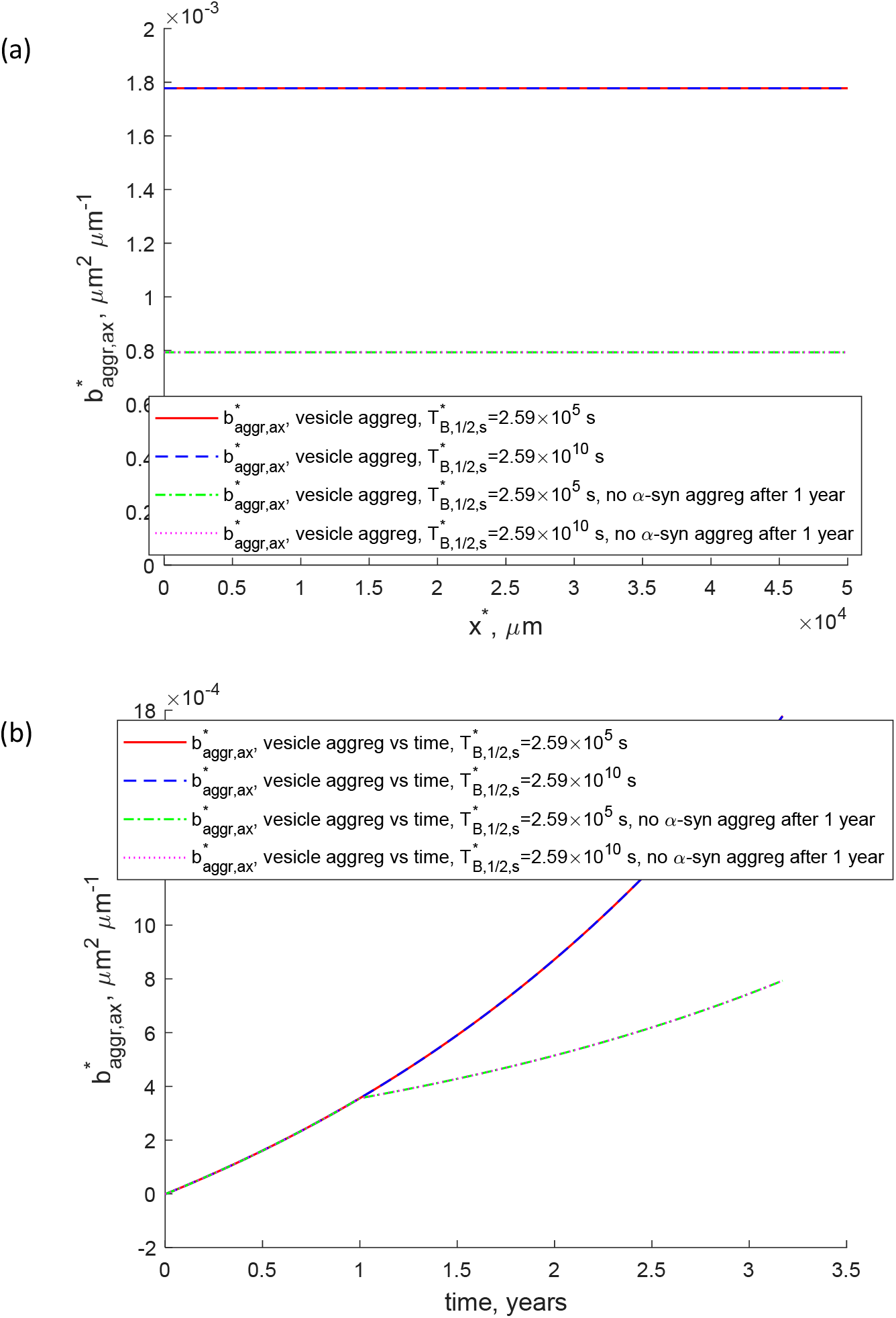
(a) Total area of the membrane in all vesicle aggregates in the axon, per unit length of the axon, vs *x*\*, at *t*\*=3.17 years. (b) Total area of the membrane in all vesicle aggregates in the axon, per unit length of the axon, vs time, at 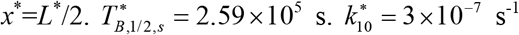.

Failure of clearance of α-syn aggregates in lysosomes may result in the accumulation of toxic oligomeric species of α-syn in the soma [75–77]. Interestingly, the excessive amount of aggregated α-syn can lead to the impairment of lysosomal activity, causing a vicious cycle of even more accumulation of misfolded α-syn in the soma [78]. To test the effects of a longer half-life of α-syn aggregates in the soma, we performed computations with two values of the half-life of α-syn aggregates in the soma: 2.59 x 10^5^ and 2.59 x 10^10^ s. The volumetric concentration of α-syn aggregates in the soma increases much faster for the larger half-life of α-syn aggregates (Fig. 8a). Since we assumed that α-syn aggregates catalyze clustering of membrane-bound vesicles in the soma (Eq. (34)), the amount of membrane contained in vesicle clusters also increases much faster for a large half-life of α-syn aggregates (Fig. 8b). Other quantities in the model are not affected (data not shown).

**Fig. 8.**
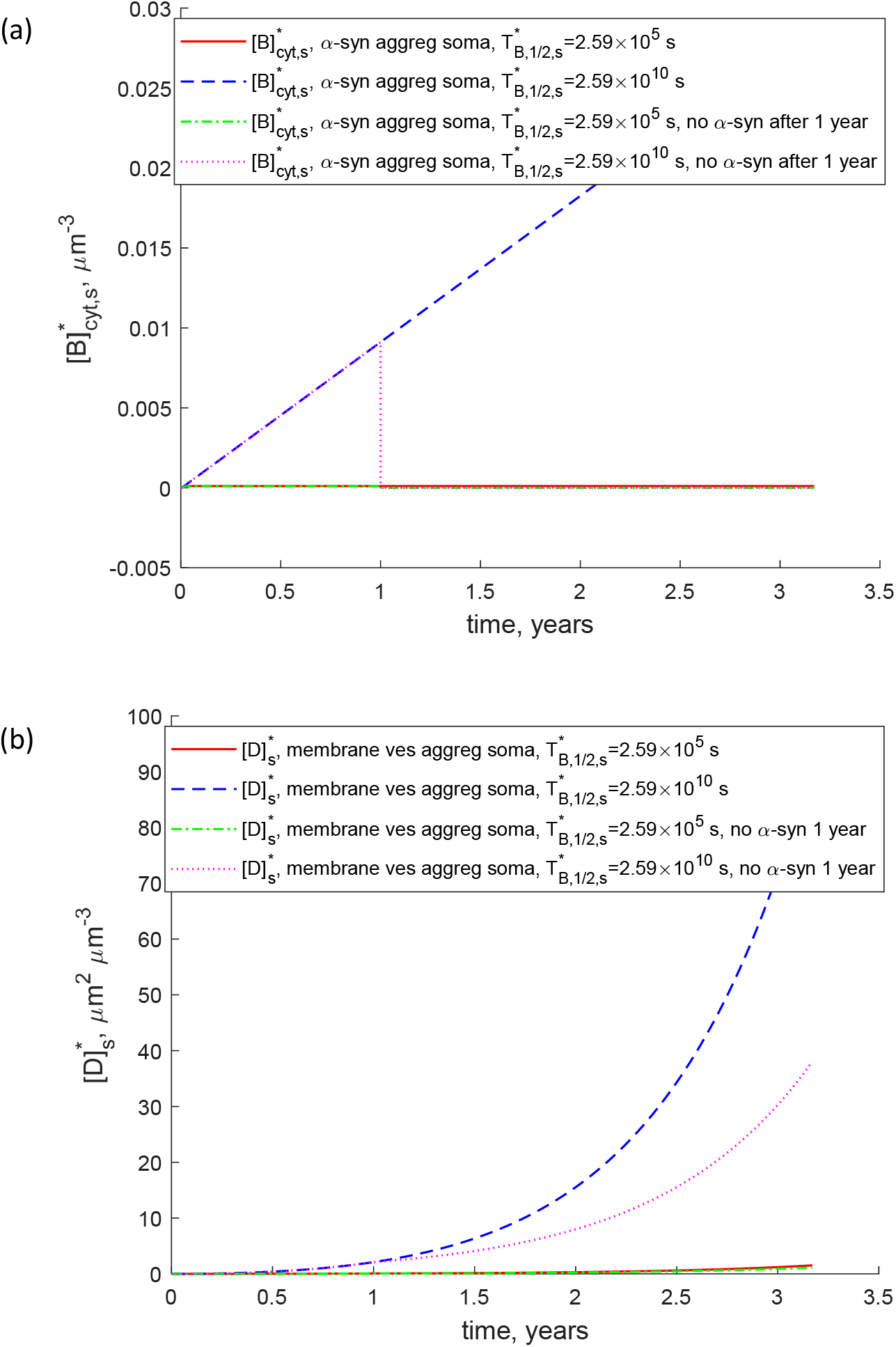
(a) Volumetric concentration of α-syn aggregates in the cytosol of the soma. (b) Volumetric concentration of membrane surrounding membrane-bound vesicles in the soma, and membrane contained in fragments and fragmented organelles in the soma. 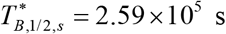 and 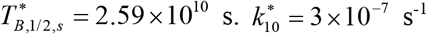.. The curves marked by “no α-syn 1 year” are computed under the assumption that all α-syn aggregates are removed (by enhancing their destruction via autophagosomes) 1 year after the beginning of the α-syn aggregation process.

It is important to understand whether the progress of PD can be stopped by removing the α-syn aggregates. The removal can potentially be accomplished by boosting autophagic or lysosomal clearance as well as using intrabodies or nanobodies for immunotherapy [79]. Another approach is using oligomer modulators, such as Anle138b, to inhibit the formation and accumulation of α-syn oligomers [80]. To simulate this situation, we removed all α-syn aggregates from the system 1 year after the formation of LBs started (dotted line in Fig. 8a). Surprisingly, the accumulation of vesicle aggregates continued, although at a slower pace (dotted line in Fig. 8b). This is explained as follows. Even if the nucleation of vesicle aggregates is suddenly stopped, the autocatalytic formation of vesicle aggregates continues. This is because in the F-W model there are two mechanisms of aggregate growth: nucleation and autocatalytic growth. This means that although PD could be initialed by α-syn oligomers, the removal of these oligomers from the brain after the formation of LBs has started may not stop the PD progression. This result is consistent with the failure of Phase III trials of Aβ-targeting drugs for Alzheimer’s disease (AD). Since the Aß deposition may begin two decades or even more before clinical symptoms of AD, the administered AD treatment in these trials may simply be too late to stop the disease [81,82].

### 3.2. Investigating sensitivity of the area of membrane contained in vesicle aggregates in the soma to model parameters

We investigated how the area of the membrane contained in vesicle aggregates by the end of the simulation (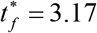 years) depends on model parameters. The analysis was performed by computing the local sensitivity coefficients, which are first-order partial derivatives of the membrane area with respect to the parameters [83–86]. The sensitivity coefficient of 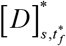 to parameter 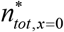, for example, was calculated as follows:

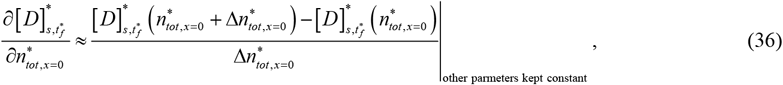

where 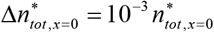. The independence of the sensitivity coefficients of the step size was tested by using various step sizes.

We then calculated non-dimensionalized relative sensitivity coefficients following [84,87] as, for example:

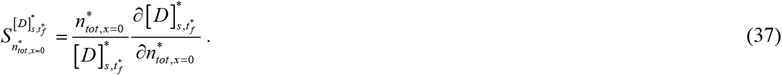

The relative sensitivity coefficients are reported in Tables 3, 5, 7, 9, and 11. The concentration of vesicle aggregates in the soma is highly sensitive to the following parameters: 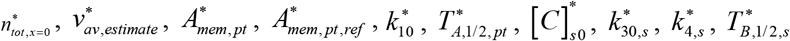, and 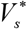 (the range of the absolute value of relative sensitivity is 0.97-4.3). Among those, 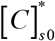, which is the volumetric concentration of membrane in the membranebound vesicles in the soma, and 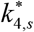, which is the rate constant for the second step of the F-W model for vesicle aggregation in the soma, produce the highest relative sensitivity. The concentration of vesicle aggregates in the soma exhibits medium sensitivity to the following parameters: 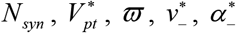, and 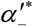 (the range of the absolute value of relative sensitivity is 5.2 x10^-4^-2.4 x10^-3^). The concentration of vesicle aggregates in the soma exhibits low sensitivity to the following parameters: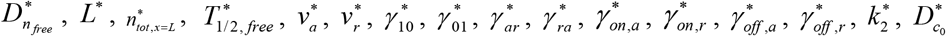, and *λ* (the range of the absolute value of relative sensitivity is 7.2 x 10^-13^ – 4.2 x 10^-5^). Finally, parameters given in Table 9 (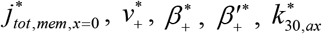, and 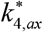) do not affect vesicle aggregates in the soma (the relative sensitivity is 0) because these parameters affect only the events occurring downstream of vesicle aggregation in the soma. However, the parameters that are given in Table 9 affect vesicle aggregation in the axon, as shown in Table S1 in the Supplemental Materials.

**Table 10.** Dependent variables in the model describing the formation of vesicle aggregates and α-syn aggregate destruction in the soma.

| Symbol | Definition | Units |
| --- | --- | --- |
| $[B]_{cyt,s}^*$ | Volumetric concentration of α-syn aggregates in the cytosol of the soma | $\mu\text{m}^{-3}$ |
| $[D]_s^*$ | Volumetric concentration of membrane in membrane fragments and fragmented organelles that were found in LBs in [5] | $\mu\text{m}^2\mu\text{m}^{-3}$ |

**Table 11.** Parameters characterizing the formation of vesicle aggregates and α-syn aggregate destruction in the soma. The sensitivity of the area of membrane contained in vesicle aggregates in the soma to these parameters (see section 3.2) is given in the last column. The cells with the largest sensitivity coefficients are shaded grey (all sensitivity coefficients given in Table 11 happen to be large).

| Symbol | Definition | Units | Value or range | Reference(s) | Value(s) used in computations | $S_{\#}^{[D]_{s,t}^*}$ |
| --- | --- | --- | --- | --- | --- | --- |
| $[C]_{s0}^*$ | Volumetric concentration of membrane in the membrane-bound vesicles in the soma | $\mu\text{m}^2\mu\text{m}^{-3}$ | 0.9 <sup>a</sup> | | 0.9 | 4.27 |
| $k_{30,s}^*$ | Proportionality constant determining the first step of the F-W model for vesicle aggregation in the soma | $\mu\text{m s}^{-1}$ | $2.78 \times 10^{-7} - 9.44 \times 10^{-5}$ | [32] | $10^{-5}$ | 1.00 |
| $\frac{k_{4,s}^* N_A^*}{3A_{s,1}}$ <sup>b</sup> | Rate constant for the second step of the F-W model for vesicle aggregation in the soma | $\mu\text{m s}^{-1} \text{ mol}^{-1}$ | $1.19 \times 10^{15} - 5.00 \times 10^{15}$ | [32] | $3 \times 10^{15}$ | 3.27<br>$(S_{k_{4,s}^*}^{[D]_{s,df}^*})$ |
| $T_{B,1/2,s}^*$ | Average half-life of α-syn aggregates before their destruction in somatic lysosomes | s | $5.76 \times 10^4 - 7.78 \times 10^5$ | [72] | $2.59 \times 10^5$ | $9.84 \times 10^{-1}$ |
| $V_s^*$ | Volume of the soma | $\mu\text{m}^3$ | $4.189 \times 10^3$ <sup>c</sup> | [73] | $4.189 \times 10^3$ | -1.00 |

<sup>a</sup> We assumed that the area of the internal membrane inside the soma is three times larger than the area of the plasma membrane.
<sup>b</sup> For the characteristic area of the plasma membrane we used $A_{s,1} = \pi(1 \mu\text{m})^2$ , which is the area of the plasma membrane around a sphere with a diameter of 1 $\mu\text{m}$ .
<sup>c</sup> Based on a representative soma diameter, $D_s^*$ , of 20 $\mu\text{m}$ .

If misfolded α-syn is the disease-causing agent, why does the premotor phase of PD span many years [3,18] while transport of misfolded α-syn aggregates from the synapse to the soma take days or weeks at most [19]? To answer this question, we compared the results presented in Fig. 8 with the results computed for a three orders of magnitude smaller rate constant for the first step of the F-W model for aggregation of α-syn monomers in the presynaptic terminal, 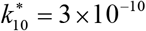. The reduction of the α-syn aggregation rate leads to a significant reduction of the concentration of α-syn in the soma (Fig. 9a) as well as to a significant reduction of the concentration of aggregates of membrane-bound vesicles in the soma (Fig. 9b), which are the main component of LBs. The slow rate of PD progression can then be explained by the need to produce α-syn aggregates first, which then catalyze the development of LBs. The formation of α-syn oligomers alone is insufficient for the development of PD and just initiates the process, which takes much longer than the time required for the production and transport of α-syn aggregates.

**Fig. 9.**
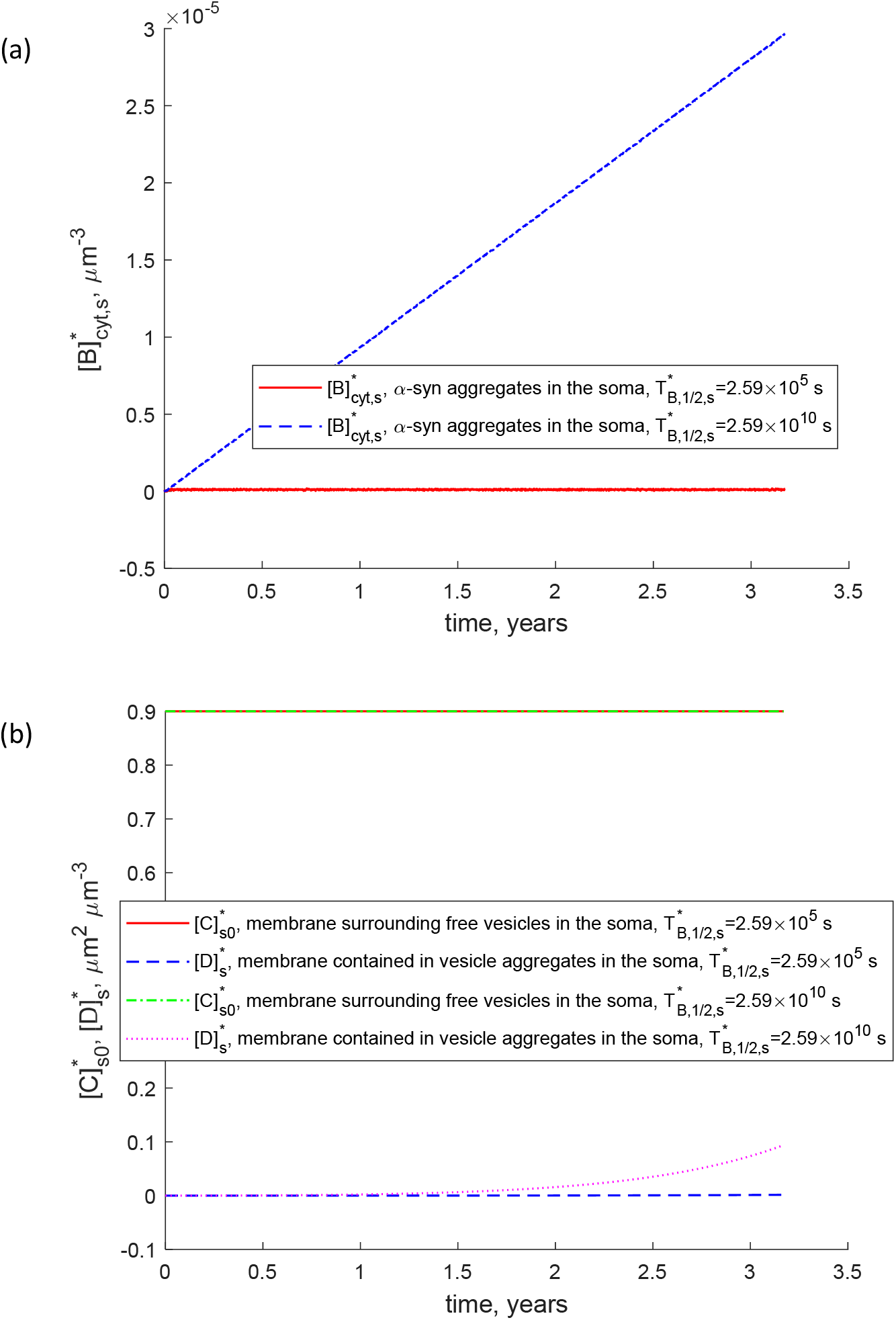
(a) Volumetric concentration of α-syn aggregates in the cytosol of the soma. (b) Volumetric concentration of membrane surrounding membrane-bound vesicles in the soma, and membrane contained in fragments and fragmented organelles in the soma. 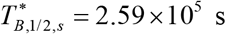 and 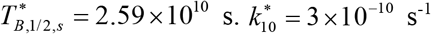.

## 4. Discussion, limitations of the model, and future directions

The purpose of this paper was to develop a minimal model for studying the possible effects of aggregated forms of α-syn on LB formation. We propose that such α-syn oligomers and fibrils work as catalysts, facilitating the formation of LBs and LNs, but not becoming a part of them.

The model simulates the transport of α-syn in the axon, production of α-syn aggregates in the synapse due to α-syn interaction with the synaptic membrane, transport of aggregates to the soma for degradation in somatic lysosomes, and catalysis of clustering of membrane-bound vesicles in the soma and the axon by α-syn aggregates.

There is still debate on whether endogenous α-syn oligomers are harmless but can convert to toxic oligomers or all α-syn oligomers are toxic [88]. The model predicts that the formation of a small amount of α-syn aggregates in the presynaptic terminal because of the folding of α-syn monomers due to their interaction with the plasma membrane is a normal process. This model prediction is in agreement with [18] that reported that misfolded α-syn is abundant in presynaptic terminals. The aggregates produced in the terminal are then transported to the soma for degradation in the lysosomes. However, on their way to lysosomes aggregates can catalyze clustering of membrane-bound vesicles. This can happen both in the axon and in the soma at any time until α-syn aggregates reach lysosomes.

Interestingly, the model predicts that α-syn monomers, transported by slow axonal transport, move against their concentration gradient, with the lowest concentration at the soma and the highest concentration at the presynaptic terminal. The distribution of α-syn monomers along the axon length is, in principle, experimentally measurable and our predictions about α-syn distribution, displayed in Fig. 3a, can be tested in future experiments. Studies of subcellular localization of α-syn have been reported, such as [89]. Notably, such studies tend to focus on the distribution of α-syn aggregates because, due to their density, they are much more obvious under light or electron microscopy. However, one can envision an experiment wherein labeling (fluorescence or otherwise) of endogenous α-syn is used to uncover the underlying monomer distribution.

According to the model, α-syn aggregates and membrane-bound vesicles, moved by fast axonal transport, exhibit a uniform concentration along the axon. Modeling results confirm that failure of clearance of oligomeric species of α-syn in somatic lysosomes (which was simulated by increasing the half-life of α-syn aggregates in the soma) may lead to the formation of LBs in the soma. According to our hypothesis, this occurs due to more catalytic activity of α-syn aggregates, which leads to the formation of vesicle aggregates in the soma. These aggregates later develop into LBs. This result is in agreement with a mouse model of PD, which shows that low levels of lysosomes precede accumulation of autophagosomes and thus suggests the importance of this clearance pathway in PD [56]. Notably, a recently published set of reporters of α-syn conformation [90] have utility for experimentally validating this phenomenon. These α-syn reporters are based upon fluorescence resonance energy transfer (FRET); changes in α-syn conformation are transduced as changes in FRET efficiency. Therefore, in principle, these reporters allow one to study, at the subcellular level and with high temporal resolution, the progression of α-syn aggregation. Hence, these reporters are broadly useful, not just for validating the previously described hypothesis, but also for experimentally confirming many of the other modeling derived hypotheses described here.

Since the formation of LBs is an autocatalytic process, and old vesicle aggregates can catalyze the formation of new vesicle aggregates even in the absence of α-syn aggregates, the model predicts that removing α-syn aggregates after the formation of LBs has begun may not stop the PD progression. Thus after a certain point, the process of LB formation can become self-sustaining. This prediction is both testable and falsifiable. This may have significant implications on PD treatment strategies, as if this prediction is true we should expect to see a broad failure of PD clinical trials focused on therapeutics which reduce the burden of α-syn aggregates after PD is already clinically diagnosed. Indeed, if true, this suggests that it is critical to begin treatment of PD early, before classical clinical signs are apparent, and hence necessitating better technologies for early detection of PD.

Future versions of the model should take into account the following. (i) Effect of a complex structure of axonal arbor. DA neurons in SNc have hundreds of thousands of synapses, which is two orders of magnitude more than other neurons in basal ganglia [52,53,91]. (ii) Simulating the possible effect of traffic jams caused by aggregation of membrane-bound vesicles. This is important because synaptic degeneration in PD may be caused by the lack of presynaptic proteins, called “vacant synapses” [92]. (iii) Simulating the effect of the 10-15% of α-syn monomers that may be transported in fast axonal transport [23]. (iv) A more precise model of α-syn aggregation in the presynaptic terminal is needed, which should include different pools of α-syn: unbound, soluble monomers, monomers bound to presynaptic vesicles, and microaggregates [93]. An α-syn polymerization model capable of distinguishing between different intermediate forms of α-syn aggregates is needed as well. This is especially important because α-syn oligomers are potentially the true disease-causing species, while large α-syn aggregates such as LBs may be neuroprotective [19,49,94]. Future research should use a combination of modeling and experimentation to answer the above-mentioned questions. (v) The concentrations shown in Fig. 4 quickly reach their steady-state values. The concentrations shown in Figs. 5, 6, and 7a are uniform along the axon length. These observations may allow, in future research, for the model to be simplified via decreasing the number of model parameters. (vi) Future work should also investigate whether aggregation of membrane fragments into Lewy bodies can be caused or facilitated by the impairment of lysosomes by α-syn aggregates [95].

## Acknowledgment

IAK acknowledges the fellowship support of the Paul and Daisy Soros Fellowship for New Americans and the NIH/National Institutes of Mental Health (NIMH) Ruth L. Kirchstein NRSA (F30 MH122076-01). AVK acknowledges the support of the National Science Foundation (award CBET-2042834) and the Alexander von Humboldt Foundation through the Humboldt Research Award.

## Supplemental Materials

**Table S1.**
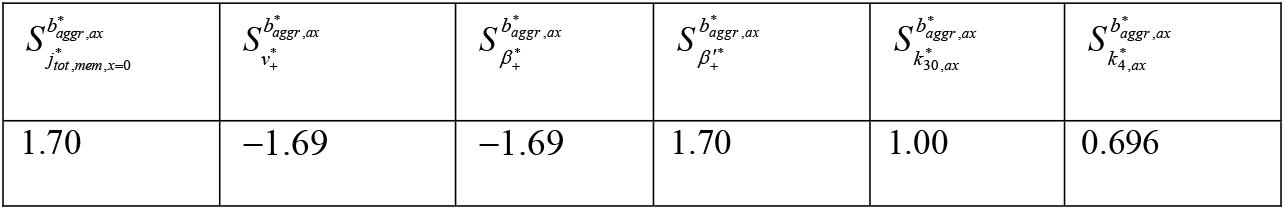
Relative sensitivity of the total area of membrane in all vesicle aggregates in the axon (that later become LNs), 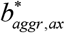, to parameters given in Table 9. Computations were performed using Eqs. (36) and (37) with (for example) 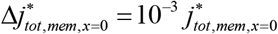.

