## Supplemental Table 1 for "Can the lack of fibrillar form of alpha-synuclein in Lewy bodies be explained by its catalytic activity?" (from another version of this preprint: version 1, 2021-05-09)

I. A. Kuznetsov[(a), (b)] and A. V. Kuznetsov[(c)]

[(a)]Perelman School of Medicine, University of Pennsylvania, Philadelphia, PA 19104, USA

[(b)]Department of Bioengineering, University of Pennsylvania, Philadelphia, PA 19104, USA

[(c)]Dept. of Mechanical and Aerospace Engineering, North Carolina State University,

Raleigh, NC 27695-7910, USA;

### Supplemental Materials

#### S1. Tables

Table S1. Relative sensitivity of the total area of membrane in all vesicle aggregates in the axon (that later become LNs), $b^*_{aggr,ax}$, to parameters given in Table 9. Computations were performed using Eqs. (36) and (37) with (for example) $\Delta j^*_{tot,mem,x=0} = 10^{-3} j^*_{tot,mem,x=0}$.

| $S^{b^*_{aggr,ax}}_{j^*_{tot,mem,x=0}}$ | $S^{b^*_{aggr,ax}}_{v^*_+}$ | $S^{b^*_{aggr,ax}}_{\beta^*_+}$ | $S^{b^*_{aggr,ax}}_{\beta'^*_+}$ | $S^{b^*_{aggr,ax}}_{k^*_{30,ax}}$ | $S^{b^*_{aggr,ax}}_{k^*_{4,ax}}$ |
|---|---|---|---|---|---|
| 1.70 | −1.69 | −1.69 | 1.70 | 1.00 | 0.696 |
